# The lysosomal carrier SLC29A3 supports anti-bacterial signaling and promotes autophagy by activating TRPML1 in mouse dendritic cells

**DOI:** 10.1101/2025.06.11.659112

**Authors:** Daniel J. Netting, Cynthia López-Haber, Zachary Hutchins, José A. Martina, Rosa Puertollano, Adriana R. Mantegazza

## Abstract

The solute carrier (SLC)29A3 exports nucleosides from lysosomes into the cytosol, maintaining solute homeostasis and providing metabolic intermediates for cellular processes. Loss-of-function mutations in SLC29A3 cause H syndrome, characterized by hyperinflammation and immunodeficiency. While dysfunctions in various cell types contribute to H syndrome and to SLC29A3 deficiency in mice, the mechanisms driving hyperinflammation and immunodeficiency are incompletely understood. Remarkably, the possible role played by dendritic cells (DCs), the most efficient antigen presenting cells and the main link between innate and adaptive immune responses, remains unknown. We show that, in murine DCs, SLC29A3 is recruited to phagosomes after bacterial capture, maintains phagosomal pH homeostasis, and ensures optimal phagosomal signaling to the production of IL-6, IL-12, and CCL-22. In addition, SLC29A3 promotes Ag presentation on MHC-II molecules to initiate adaptive immune responses. Notably, SLC29A3 supports the activity of the lysosomal calcium channel TRPML1, promoting transcription factor TFEB nuclear translocation and inducing autophagy, a major anti-inflammatory mechanism. Overexpression of human SLC29A3, but not the transport mutant G437R, in SLC29A3-deficient murine DCs restores cytokine production in response to bacteria phagocytosis, suggesting that SLC29A3 transport activity is required to drive anti-bacterial phagosomal signaling. Our data indicate that SLC29A3 plays a dual role in supporting immune function in DCs by promoting effective anti-microbial signaling and Ag presentation and inducing autophagy to control inflammation. Our findings also uncover a novel TRPML1-dependent mechanism by which SLC29A3 activates TFEB and suggest that defects in phagosomal signaling, TFEB activation and autophagy may contribute to immunodeficiency and hyperinflammation in SLC29A3 disorders.

## Introduction

The importance of maintaining proper solute or nutrient transport to ensure cell and immune fitness is evidenced by the association between dysfunctions in many <u>s</u>olute <u>c</u>arrier transporters (SLCs) and human metabolic and inflammatory disorders. These include lymphadenopathy-histiocytosis plus syndrome or H syndrome – recently incorporated into the families of immunodeficiencies or “inborn errors of immunity” and “inborn errors of metabolism”(1-5). H syndrome is caused by loss-of-function mutations in the lysosomal nucleoside transporter SLC29A3, the only known intracellular nucleoside transporter(6-8). H syndrome encompasses a wide range of symptoms, including histiocytosis, which is characterized by the accumulation of dysfunctional phagocytes(4, 9-11). While contributions from various cell types have been described for H syndrome and SLC29A3 deficiency in humans and mice(12-15), the molecular mechanisms linking SLC29A3 dysfunction to hyperinflammation and immunodeficiency have not been completely elucidated. Furthermore, whether and how SLC29A3 regulates the function of dendritic cells (DCs), the main cellular link between innate and adaptive immunity(16, 17), to promote anti-bacterial immune responses, remains unknown.

Anti-microbial functions in phagocytes such as macrophages (MΦs) and DCs, begin with the enclosure of captured bacteria into newly formed phagosomes(18-20). Phagosomes subsequently mature through interactions with the endolysosomal system, acquiring endosomal <u>p</u>attern recognition receptors (PRRs) like <u>T</u>oll-like receptors (TLRs, which sense microbial components and signal to anti-microbial cytokine pathways and Ag presentation) and lysosomal degradative enzymes(20-31). We and others have shown that some lysosomal SLCs are also recruited to phagosomes during maturation(32-34). Phagolysosomal SLCs may contribute to phagosome and cell homeostasis by exporting solutes into the cytosol, providing building blocks for anabolic cellular processes and maintaining optimal membrane tension(35, 36). In addition, some lysosomal SLCs, like SLC15A4, link nutrient transport and sensing to anti-bacterial pro-inflammatory responses(32, 37).

In MΦs, SLC29A3 loss of function is associated with increased levels of phagolysosomal nucleosides and nucleobases, which impairs phagosomal acidification and bacterial clearance(13). This may also affect the generation of <u>p</u>athogen-<u>a</u>ssociated <u>m</u>olecular <u>p</u>atterns (PAMPs) for recognition by phagosomal TLRs(38). Therefore, we hypothesized that nucleoside accumulation in SLC29A3-deficient (SLC29A3^-/-^) DC phagosomes after bacteria phagocytosis may increase phagosomal pH and impair phagosomal TLR signaling pathways, dampening cytokine and chemokine production and negatively impacting Ag presentation, a major feature in DCs. This scenario would necessitate additional phagocyte recruitment to control bacterial infections *in vivo*, which could contribute to histiocytosis and inflammation(39).

Moreover, we previously showed that the lysosomal transporter SLC15A4, associated with inflammatory disorders(40, 41), promotes inflammatory responses in DCs by conveying nutrient availability to the master regulator of cell fitness, <u>m</u>echanistic target <u>o</u>f rapamycin <u>c</u>omplex 1 (mTORC1)(42, 43) and thereby inhibiting autophagy, a major catabolic and anti-inflammatory process(32, 44). Thus, we hypothesized that hyperinflammation in H syndrome might also be associated with dysregulated autophagy, and that defects in SLC29A3 function could impact mTORC1 function and/or the activity of transcription factor <u>EB</u> (TFEB), the master regulator of lysosomal function and autophagy, after phagocytosis(45, 46). Indeed, after phagocytosis in MΦs, calcium efflux from phagosomes, mainly mediated by the lysosomal transient receptor <u>p</u>otential <u>m</u>ucolipin-<u>1</u> (TRPML1), drives TFEB nuclear translocation via the calcium-dependent phosphatase calcineurin, leading to the transcription of the coordinated lysosomal expression and regulation gene network(47-51). TFEB has been traditionally linked to nutrient sensing, metabolism, lysosomal biogenesis, and autophagy, but it is also linked to anti-microbial cytokine responses in MΦs, like the transcription factors NF-κB and AP-1(48, 52, 53). Notably, lysosomal adenosine accumulation due to adenosine deaminase deficiency and increased pH are shown to impair TRPML1 activity(54-57). Therefore, we hypothesized that adenosine accumulation and potentially increased phagosomal pH in SLC29A3-deficient DCs may impair both phagosomal TLR signaling and TFEB nuclear translocation, affecting cytokine/ chemokine anti-microbial programs and autophagy in DCs.

We show herein that SLC29A3 is recruited to DC phagosomes during maturation and supports phagosomal acidification. In addition, SLC29A3 promotes TLR/NF-κB-dependent IL-6, IL-12, and CCL22 production after Salmonella *enterica* serovar typhimurium (STm) phagocytosis by bone marrow-derived and tissue-resident splenic DCs. Furthermore, SLC29A3 is required by DCs for optimal MHC-II presentation to CD4+ T cells to initiate adaptive immune responses. Mechanistically, we show that SLC29A3 supports TRPML1 function on phagolysosomes and promotes TFEB nuclear translocation, sustaining autophagy and likely anti-microbial programs. Altogether, our data uncover previously neglected molecular mechanisms governing SLC29A3-associated inflammation and immune responses in DCs, key players in the orchestration of pro-inflammatory responses upon phagocytosis.

## Results

### SLC29A3 is recruited to phagosomes and promotes phagosomal tubulation and acidification in DCs

SLC29A3 bears an N-terminal dileucine lysosomal targeting motif and is, therefore, located on lysosomal membranes(6, 11). To confirm the lysosomal localization of a GFP-tagged human SLC29A3 construct (GFP-SLC29A3), we pulsed BMDCs expressing GFP-SLC29A3 with the fluorescent dye LysoTracker Red to identify acidic organelles in live cells. As expected, GFP-SLC29A3 was detected on lysosomal membranes, surrounding LysoTracker Red positive vesicles in BMDCs (**Fig. 1A**). We then investigated whether SLC29A3 is recruited to phagosomes and phagosomal tubules after phagocytosis. Previously, we showed that DC phagosomes emit tubules during their maturation process by a mechanism dependent on TLR signaling, to support Ag presentation to T cells(58). Increased phagosomal surface may also favor solute export to the cytosol(37). We found that GFP-SLC29A3 was recruited to BMDC phagosomes and phagosomal tubules 2 h after phagocytosis of Texas red (TxR)-conjugated OVA and LPS (to stimulate TLR4)-coated polystyrene beads (TxR-OVA/LPS beads; **Fig. 1B**). Furthermore, by proteomic analyses on isolated mature phagosomes from BMDCs pulsed with LPS-coated beads and chased for 2 h, we detected peptides from SLC29A3 (**Fig. 1C**), confirming previous reports(34). Notably, SLC29A3^-/-^ BMDC phagosomes are impaired in their ability to form phagolysosomal tubules compared to WT (**Fig. 1D, E**), in BMDCs pre-incubated with apoptotic splenocytes as a source of nucleosides. This suggests that SLC29A3 regulation of phagosomal membrane tension or deformation(35) and/or phagosomal TLR signaling(58) may promote phagosomal membrane dynamics.

**Figure 1.**
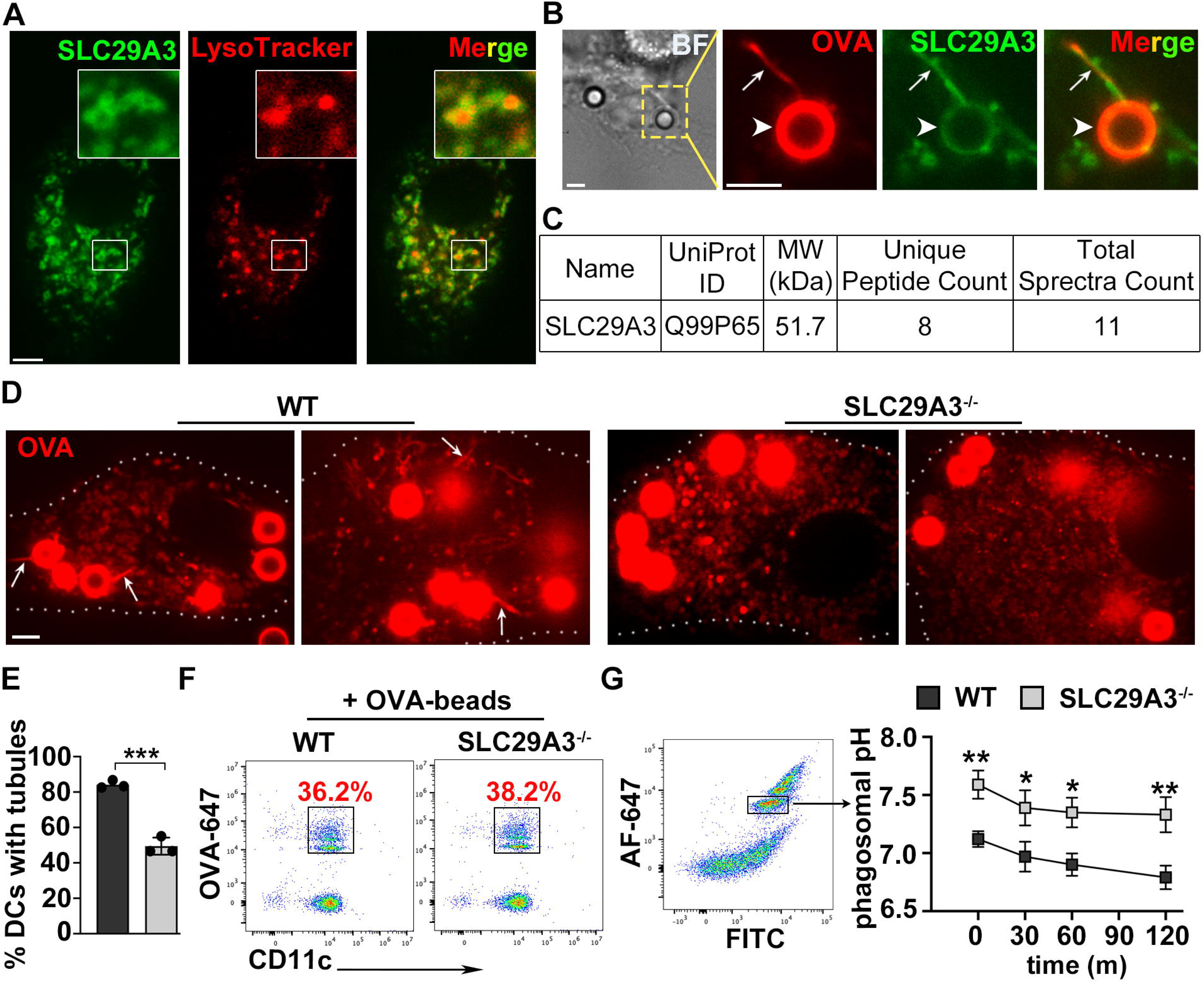
SLC29A3 is recruited to phagosomes and promotes phagosomal tubulation and acidification in DCs. WT BMDCs were transduced with GFP-SLC29A3 (A, B) or WT and SLC29A3^-/-^ BMDCs or splenic DCs were non-transduced (C-G). **A.** Cells were analyzed 1 hour after treatment with LysoTracker Red by live-cell imaging. Representative images of three independent experiments. **B, D, E.** GFP-SLC29A3-expressing WT (B) and non-transduced WT and SLC29A3^-/-^ BMDCs (D, E) were pulsed with TxR-OVA/LPS beads and analyzed 2 h after the pulse by live-cell imaging. Representative images of three independent experiments. Quantification of the percent of BMDCs showing phagosomal tubules in 100 cells, in 3 independent experiments (E). **C.** WT BMDCs were pulsed with magnetic LPS-beads. Isolated BMDC phagosomes 2 h after the pulse were subjected to proteomic analyses. Number of unique and total identified peptides from SLC29A3 protein is indicated. **F.** WT and SLC29A3^-/-^ splenic DCs were pulsed with OVA-AF647 beads. Phagocytic capacity was measured by flow cytometry. Percent of DCs that phagocytosed beads are shown in red. **G.** WT and SLC29A3^-/-^ splenic DCs were pulsed with FITC/AF647 beads to assess phagosomal pH. A population containing a similar minimal number of beads was selected as shown (*left*). Phagosomal pH was measured by flow cytometry at the indicated time points. pH was calculated by measuring the ratio between FITC and AF647 MFIs and interpolating these values into a standard curve of fixed pH values (*right*). Data represent mean ± SD. * p<0.05; ** p<0.01. Two-tailed unpaired Student’s T-test. *Arrows*, phagosomes; *arrowheads*, phagosomal tubules. Scale bar, 3μm.

SLC29A3 supports phagosomal acidification in MΦs(13). However, given the differential phagosomal properties between MΦs and DCs(59, 60), we assessed whether SLC29A3 played a similar role in DCs. To this end, we isolated tissue-resident splenic DCs, which accumulate nucleosides and nucleobases *in vivo*. While differences in the relative frequencies of monocytes and T cells were reported in SLC29A3^-/-^ mice(12, 14, 15), SLC29A3 deficiency did not affect the differentiation of splenic DCs (**Supp. Fig. 1A**). Isolated splenic DCs from wild-type (WT) and SLC29A3^-/-^ mice were pulsed with fluorescein isothiocyanate (FITC; pH sensitive) and Alexa Fluor 647 (AF-647; pH insensitive)-coated beads and phagosomal pH was measured over time after phagocytosis by flow cytometry. Phagocytic capacity was similar between WT and SLC29A3^-/-^ DCs (**Fig. 1F**). We found that even though phagosomal pH decreases mildly in DCs compared to macrophages, as we and others previously described(61, 62), phagosomes in SLC29A3^-/-^ DCs are significantly more alkaline than their WT counterparts (**Fig. 1G**), suggesting that SLC29A3 is required for proper phagosomal acidification after phagocytosis in DCs.

### SLC29A3 promotes cytokine and chemokine production after bacteria phagocytosis

Given that phagosomal pH was more alkaline in SLC29A3^-/-^ splenic DCs (**Fig. 1G**), and that acidification is required for phagosomal content degradation and further exposure of PAMPs for recognition by phagosomal PRRs, we investigated whether phagosomal pro-inflammatory signaling was affected by SLC29A3 after bacteria phagocytosis in DCs. Similarly to splenic DCs, SLC29A3 deficiency did not affect BMDC differentiation, maturation or phagocytic capacity **(Supp. Fig. 1B-D**). WT and SLC29A3^-/-^ BMDCs and splenic DCs were stimulated with gentamicin-killed STm (STm Gm) or live-attenuated STm lacking flagella (StmΔfla), to preserve cell viability(32) and stimulate PRRs, at a multiplicity of infection (MOI) of 3 for 3 h. We found that the secretion of IL-6, IL-12, and CCL-22 was significantly impaired in SLC29A3^-/-^ BMDCs compared to WT (**Fig 2A, C, E**), suggesting that SLC29A3 is required for optimal cytokine and chemokine secretion. We also detected reduced IL-6, and more significantly reduced IL-12 and CCL22 mRNA levels both by qPCR analyses and RNA seq studies, indicating impaired transcriptional responses in SLC29A3^-/-^ DCs (**Supp. Fig. 2A-C**). Cytokine secretion was even more severely impaired in SLC29A3^-/-^ tissue resident splenic DCs, likely due to nucleoside accumulation *in vivo* (**Fig. 2B, D, F**).

**Figure 2.**
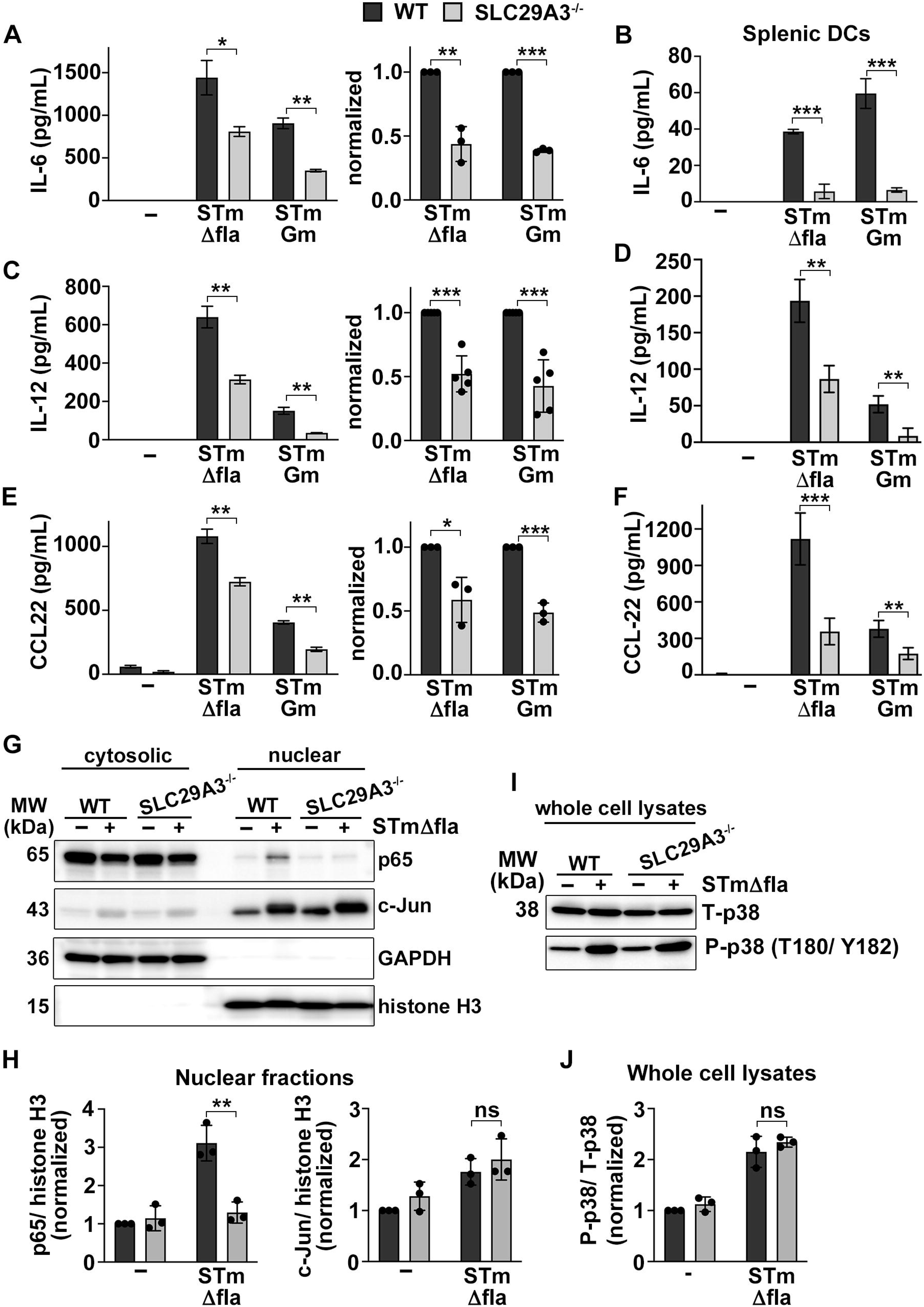
SLC29A3 promotes cytokine and chemokine production by DCs after phagocytosis. WT and SLC29A3^-/-^ BMDCs (A, C, E, G-J) were stimulated with STmΔfla or Gm-killed STm at MOI=3 (A, C, E) or MOI=30 (G-J). WT and SLC29A3-/- splenic DCs were stimulated with STmΔfla or Gm-killed STm at MOI=10 (B, D, F). **A-F.** Cell supernatants collected 3 h after stimulation were assayed for IL-6 (A, B), IL-12 (C, D), and CCL22 (E, F) by ELISA. **A, C, E**. Representative experiments (*left*). Average values of three independent experiments were normalized to WT BMDCs (*right*). Data represent mean ± SD. **B, D, F.** Representative plots of 3 independent experiments. Data represent mean ± SD. **G, H.** Subcellular fractionations were performed 3 h after stimulation. Nuclear and cytosolic fractions were immunoblotted for p65, c-Jun, GAPDH, and histone-H3. GAPDH and histone H3 are controls for cytosolic and nuclear fractions, respectively. Representative immunoblots of three independent experiments (G). Quantification of three independent experiments was normalized to untreated WT (H). Data represent mean ± SD. **I, J.** Whole-cell lysates were prepared 3 h after stimulation and immunoblotted for total (T) and phospho (P) p38. Representative immunoblots of three independent experiments (I). Quantification of three independent experiments was normalized to untreated WT (J). Data represent mean ± SD. * p<0.05; ** p<0.01; *** p<0.001. Two-tailed unpaired Student’s T-test.

To investigate the phagosomal signaling pathways leading to cytokine secretion regulated by SLC29A3, we stimulated BMDCs with STmΔfla and performed subcellular fractionations. We found that the translocation to the nucleus of the NF-κB subunit p65 was reduced in SLC29A3^-/-^ BMDCs compared to WT, suggesting that the activation of NF-κB is impaired by the absence of SLC29A3. In contrast, the nuclear translocation of c-Jun, a component of the AP-1 transcription factor, was not affected, suggesting that AP-1 activation is not modulated by SLC29A3 (**Fig. 2G, H**). Consistent with this, phosphorylation of p38, a MAPK upstream of AP-1, was not affected by SLC29A3 (**Fig. 2I, J**). Impaired phagosomal signaling to NF-κB may also account at least partially for the reduced phagosomal tubulation in SLC29A3^-/-^ DCs (**Fig. 1D, E**), as we have shown in DCs with impaired TLR signaling(58, 63).

To investigate whether SLC29A3 transport function was required for the promotion of cytokine production, we transduced WT and SLC29A3 BMDCs with GFP-SLC29A3 or GFP-SLC29A3^G437R^ (G437R, a mutation detected in H syndrome patients, with impaired transport activity but conserved lysosomal localization(8, 54, 64)). Expression of these constructs was similar between cell types (**Supp. Fig. 3A**). Notably, the expression of WT human SLC29A3, but not G437R rescued the defect in cytokine secretion in SLC29A3^-/-^ BMDCs, suggesting that nucleoside transport, and not a scaffolding function, is required for cytokine production (**Fig. 3A, B**).

**Figure 3.**
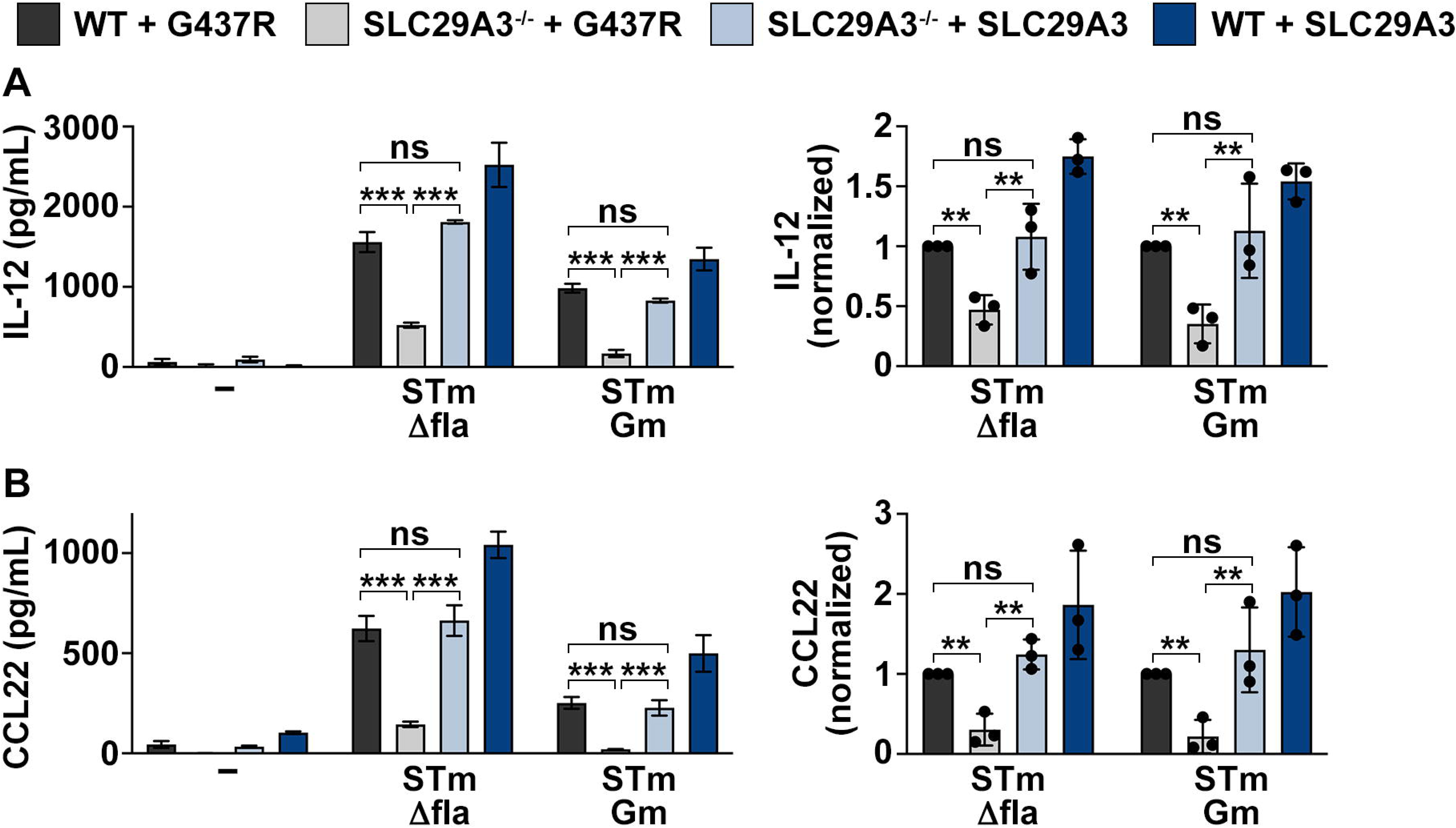
Expression of human SLC29A3, but not the transport mutant G437R restores cytokine production in SLC29A3^-/-^ DCs. WT and SLC29A3^-/-^ BMDCs or WT and SLC29A3^-/-^ BMDCs expressing GFP-SLC29A3 or GFP-SLC29A3^G437R^ were stimulated with STmΔfla or Gm-killed STm at MOI=3. **A, B.** Cell supernatants collected 3 h after stimulation were assayed for IL-12 (A) and CCL22 (B) by ELISA. Representative plots are shown (*left*). Average values of three independent experiments were normalized to WT BMDCs expressing GFP-SLC29A3^G437R^ (*right*). Data represent mean ± SD. ** p<0.01; *** p<0.001. Two-tailed unpaired Student’s T-test.

### SLC29A3 is required for optimal antigen MHC-II presentation

Considering the reduced formation of phagosomal tubules as well as the defects in phagosomal acidification in SLC29A3^-/-^ BMDCs, we investigated whether SLC29A3 was required for efficient presentation of phagocytosed or endocytosed Ags on MHC-II molecules. We pulsed splenic DCs with polystyrene beads coated with different concentrations of OVA and BSA (control), soluble OVA or the OVA peptide 323-339 (ISQAVHAAHAEINEAGR), which is presented on MHC-II molecules. We then co-cultured pulsed DCs with the CD4+ T cell hybridoma OT-IIZ, which expresses a T cell receptor that recognizes the OVA peptide 323-339 presented by DCs on MHC-II molecules(65). OT-IIZ expresses β-galactosidase (lacZ) when T cells activate in response to MHC-II:peptide recognition. We found that T cell activation in response to the presentation of the OVA-peptide, which does not need antigenic processing, was similar in co-cultures with WT or SLC29A3^-/-^ DCs. However, T cell activation in response to DCs pulsed with either OVA:BSA beads or soluble OVA was significantly impaired after co-culture with SLC29A3^-/-^ DCs compared to WT (**Fig. 4A**). Similarly, IL-2 secretion by T cells was significantly reduced after co-culture of SLC29A3^-/-^ DCs when DCs were treated with particulate or soluble OVA, but not in the case of OVA peptide (**Fig. 4B**). The expression of MHC-II molecules was similar between cell types (**Fig. 4C**), suggesting that SLC29A3 is required for efficient Ag presentation on MHC-II molecules after phagocytosis or endocytosis.

**Figure 4.**
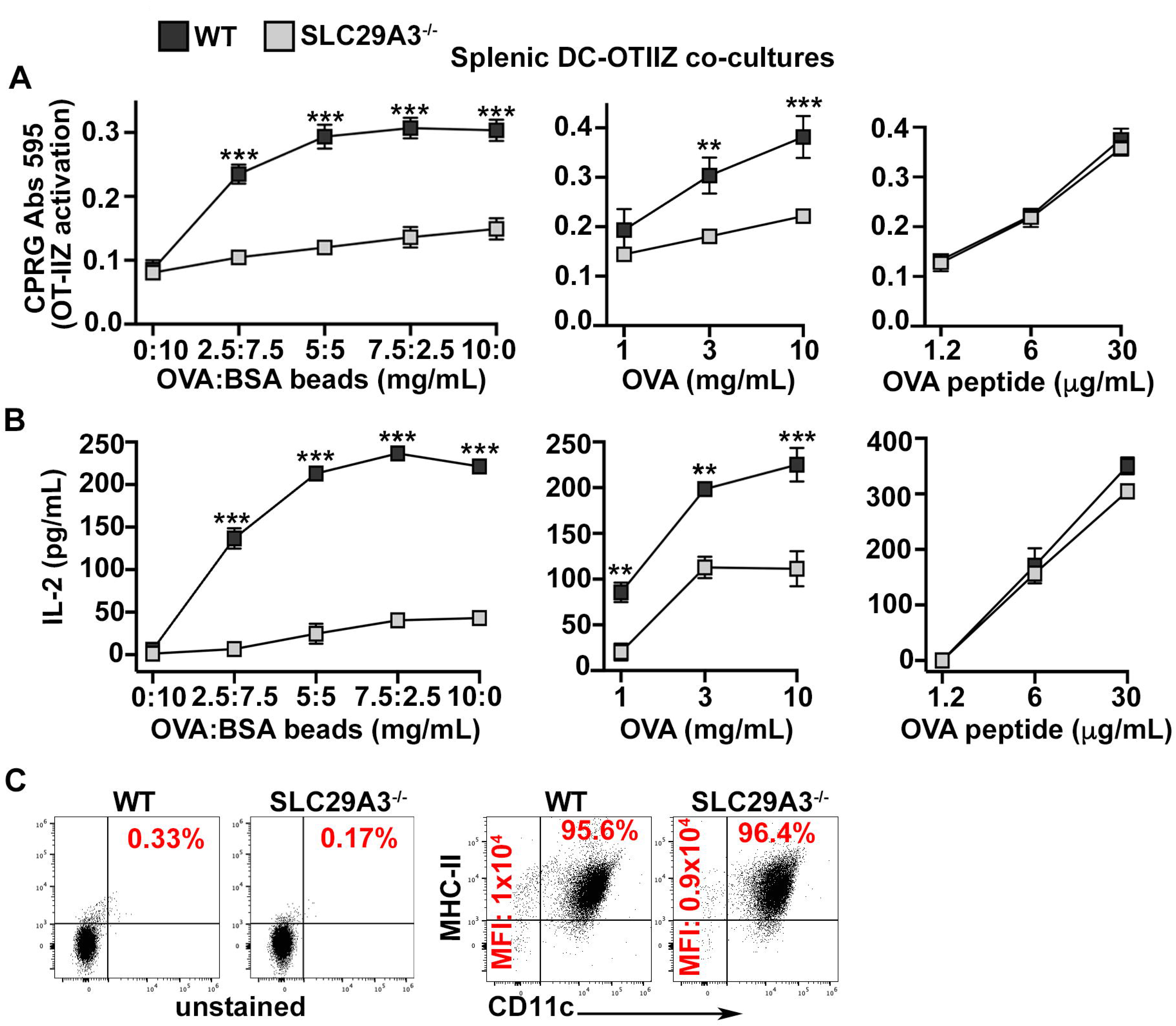
SLC29A3 is required for optimal antigen MHC-II presentation. WT and SLC29A3^-/-^ splenic DCs were pulsed with OVA:BSA coated beads, soluble OVA or OVA peptide 323-339 at the indicated concentrations for 3 h, and co-cultured with OT-IIZ hybridoma for 18 h. **A.** OT-IIZ activation was detected by adding the lacZ substrate CPRG and measuring absorbance at 595 nM after 4 h at 37°C. **B.** IL-2 secretion on DC-OT-IIZ co-culture supernatants was measured by ELISA. **C.** Expression of MHC-II molecules on isolated splenic DCs from WT and SLC29A3^-/-^ mice was measured by flow cytometry. Percent of CD11c^+^/ MHC-II^+^ DCs and MFI values are shown in red.

### SLC29A3 promotes TFEB nuclear translocation and autophagy after bacteria phagocytosis

Bacteria phagocytosis in MΦs leads to the activation of TFEB, the master regulator of lysosomal function and autophagy, but also an activator of anti-microbial cytokine responses, playing a complementary role to NF-κB(48, 53). In addition, TFEB activity was shown to promote MHC-II presentation of exogenous Ags by DCs(66). We therefore investigated whether TFEB activation was modulated by SLC29A3. At steady-state, TFEB remains phosphorylated in the cytosol, mainly by mTORC1 activity(67). After bacteria phagocytosis, TFEB translocates to the nucleus following calcineurin-mediated dephosphorylation(67-69). We stimulated human TFEB-GFP (hTFEB-GFP) expressing WT and SLC29A3^-/-^ BMDCs with STmΔfla for 3 h and assessed TFEB nuclear translocation by fluorescence microscopy and subcellular fractionation. The transduction efficiency was similar between WT and SLC29A3^-/-^ BMDCs (**Supp. Fig. 3B**). hTFEB-GFP translocated to the nucleus in WT BMDCs after bacteria phagocytosis, as observed in MΦs. In contrast, and notably, hTFEB-GFP remained largely cytosolic in stimulated SLC29A3^-/-^ BMDCs, suggesting that SLC29A3 is required for proper TFEB activation (**Fig. 5A, B**). In addition, after subcellular fractionation, hTFEB remained mostly in cytosolic fractions in SLC29A3^-/-^ BMDCs after STmΔfla stimulation, compared to WT cells, in which hTFEB was mostly detected in nuclear fractions (**Fig. 5C, E**). Also, we noted a clear shift in hTFEB molecular weight in WT cytosolic fractions, suggesting TFEB dephosphorylation, an indicator of TFEB activation and nuclear translocation(67, 68) (**Fig. 5C**, *cytosolic*). Indeed, Ser 211 phosphorylation in hTFEB is significantly reduced in WT compared to SLC29A3^-/-^ BMDCs after STmΔfla stimulation (**Fig. 5F)**. In agreement with these observations, the nuclear translocation of endogenous TFEB (mouse TFEB) was also impaired in stimulated SLC29A3^-/-^ BMDCs compared to WT (**Fig. 5D, E**). BMDC treatment with the mTORC1 inhibitor torin, to prevent TFEB phosphorylation and cytosolic retention, drove TFEB to the nucleus in both WT and SLC29A3^-/-^ BMDCs, indicating that the translocation itself is not impaired in SLC29A3^-/-^ BMDCs, and highlighting the requirement for SLC29A3 at an upstream step **(Supp. Fig. 4**).

**Figure 5.**
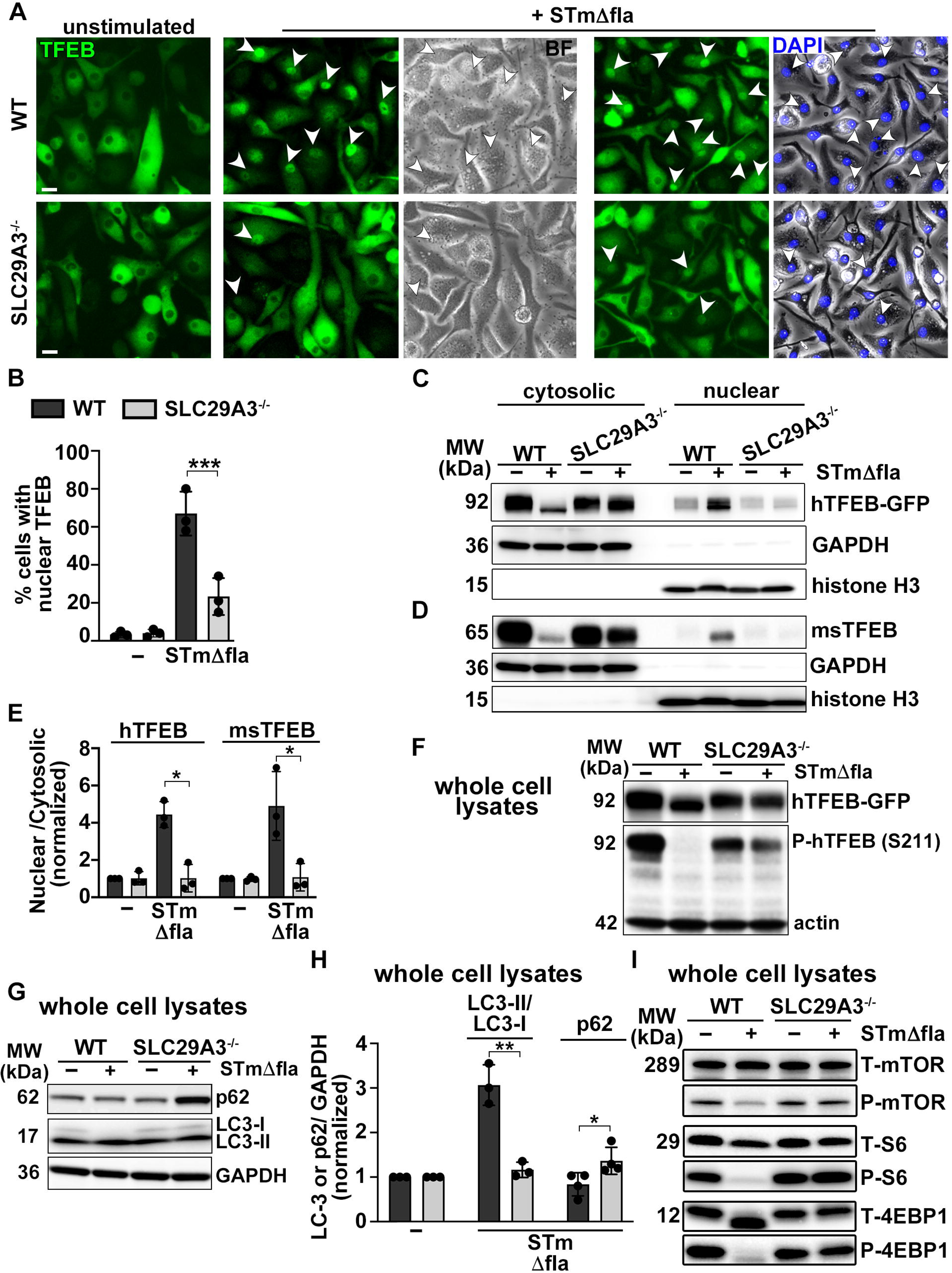
SLC29A3 promotes TFEB nuclear translocation and autophagy after bacteria phagocytosis. WT and SLC29A3^-/-^ BMDCs expressing hTFEB-GFP (A-F) or non-transduced WT and SLC29A3^-/-^ BMDCs (G-I), were unstimulated or treated with STmΔfla at MOI=30 for 3 h. **A.** TFEB-GFP nuclear translocation was assessed by fluorescence microscopy. Representative images of 3 independent experiments. *Arrows*, nuclear translocation. Magnification, 20X. Scale bars, 40 μm. **B.** Percent of cells with nuclear TFEB was quantified in a total of 300 cells in three independent experiments. **C, D.** Subcellular fractionations were performed. Nuclear and cytosolic fractions were immunoblotted for hTFEB, msTFEB, GAPDH and histone H3. GAPDH and histone H3 are controls for cytosolic and nuclear fractions, respectively. Representative immunoblots of three independent experiments. **E.** Quantification of three independent experiments was normalized to untreated WT. **F, G.** Whole-cell lysates were collected 3 h after stimulation and immunoblotted for hTFEB, phospho-hTFEB (S211) and actin (F), and LC3, p62 and GAPDH (G). **H.** Quantification of the ratio between LC3-II, LC3-I and GAPDH, and p62 and GAPDH in three independent experiments was normalized to untreated cells. **I.** Whole-cell lysates were collected 3 h after stimulation and immunoblotted for total (T) and phospho (P) mTOR, S6 and 4EBP1. Representative immunoblots of three independent experiments. Data represent mean ± SD. * p<0.05; ** p<0.01; *** p<0.001. Two-tailed unpaired Student’s T-test.

Given the role played by TFEB in the regulation of autophagy, a major catabolic and anti-inflammatory process(44), we investigated whether autophagy induction and/or flux was modulated by SLC29A3 in STmΔfla-stimulated BMDCs. We observed that autophagy induction, measured by the ratio between lipidated LC3 (LC3-II; present in autophagosomes) and unlipidated LC3 (LC3-I; cytosolic), as well as autophagic flux (measured by the degradation of p62, which links ubiquitylated substrates to LC3) were reduced in STmΔfla-treated SLC29A3^-/-^ BMDCs compared to WT (**Fig 5G**). Indeed, the LC3-II/LC3-I ratio was reduced in SLC29A3^-/-^ BMDCs compared to WT, and there was a significant accumulation of p62 in SLC29A3^-/-^ BMDCs, as detected by immunoblotting (**Fig. 5G, H**). This suggests a defective p62 degradation in the absence of SLC29A3, possibly due to elevated lysosomal pH.

Given that autophagy initiation is impaired by the phosphorylation of mTORC1(70, 71), and that the lysosomal accumulation of adenosine in SLC29A3^-/-^ hematopoietic stem cells was shown to affect the AMPK/mTORC1 axis(12), leading to mTORC1 activation, we investigated mTORC1 activity in WT and SLC29A3^-/-^ BMDCs after STmΔfla phagocytosis. We found that the phosphorylation of mTORC1 in Ser 2481 – the autophosphorylation site associated with mTORC1 catalytic activity(72) – as well as the phosphorylation of the mTORC1 substrates S6 and 4EBP1 were reduced in WT BMDCs 3 h after STmΔfla treatment, in line with the observed autophagy induction. In contrast, in SLC29A3^-/-^ BMDCs, mTORC1 and its downstream substrates remained phosphorylated in response to bacteria stimulation (**Fig. 5I**).

Altogether, our data suggest that both mTORC1 and TFEB activation are regulated by SLC29A3 to promote autophagy. In agreement with these observations, the transcriptional regulation of autophagy pathways, as well as anti-bacterial responses, was significantly different between WT and splenic DCs treated with STmΔfla, as determined by RNA seq analyses (**Supp. Fig. 5**).

### SLC29A3 promotes the activity of the lysosomal Ca^2+^ channel TRPML1 on phagolysosomes

Calcium signaling processes were also differentially regulated at transcriptional levels between WT and SLC29A3^-/-^ splenic DCs (**Supp. Fig. 5**). Activation of TFEB in response to phagocytosis is largely mediated by the Ca^2+^ channel TRPML1(51). Notably, TRPML1 activity is promoted by the acidic lysosomal pH and impaired by the lysosomal accumulation of adenosine(54). Considering the role of SLC29A3 in exporting nucleosides to the cytosol and our observations that SLC29A3-deficiency impairs phagosomal acidification in DCs (**Fig. 1G**), we hypothesized that SLC29A3 may promote TRPML1 activity in phagolysosomes by maintaining nucleoside homeostasis and supporting phagosomal acidification. To test this, we transduced BMDCs with TRPML1 fused to a genetically encoded Ca^2+^ indicator for optical imaging (GECO). TRPML1-GECO localizes to lysosomes and fluoresces in response to Ca^2+^ efflux, allowing the visualization of TRPML1-dependent Ca^2+^ dynamics after phagocytosis(54). WT and SLC29A3^-/-^ BMDCs expressing TRPML1-GECO were stimulated with gentamycin-killed STm-mCherry (to detect phagosomes) and visualized by live cell imaging 1-2 h after phagocytosis. BMDCs were pre-treated with the cell permeant reagent BAPTA-AM to chelate cytosolic calcium and imaged in calcium-free media to favor the detection of phagolysosomal Ca^2+^. We measured TRPML1 activity as the number of Ca^2+^-induced flashes of fluorescence (“hotspots”) and we differentiated between hotspots on phagosomes (periphagosomal) or away from phagosomes (lysosomal), as described(73). We found that BMDCs showed similar numbers of total Ca^2+^ hotspots (likely because lysosomes were not pre-loaded with nucleosides). However, SLC29A3^-/-^ BMDCs exhibited a significantly reduced number of periphagosomal hotspots after STm phagocytosis (**Fig. 6A, D**). To confirm that the visualized hotspots resulted from TRPML1 activity, BMDCs were pre-treated with the TRPML1 inhibitor ML-SI3 before bacteria stimulation(74). As expected, no hotspots were visualized when TRPML1 was inhibited (**Fig. 6B**). Reduced phagosomal TRPML1 activity in SLC29A3^-/-^ BMDCs was not due to reduced lysosome-phagosome interactions, as these were not impaired in SLC29A3^-/-^ BMDCs (**Fig. 6C**). In addition, SLC29A3 did not affect cytosolic calcium dynamics and/or calcium entry early after stimulation, as visualized with the fluorescent ratiometric genetically encoded cytosolic Ca^2+^ indicator GCamp6f linked to tdTomato (Salsa6f). Indeed, both Salsa6f-transduced WT and SLC29A3^-/-^ BMDCs exhibited bursts of Ca^2+^ in the cytosol over time immediately following stimulation with bacteria or LPS-beads, suggesting that cytosolic Ca^2+^ fluxes (likely resulting from Ca^2+^ release from the endoplasmic reticulum or Ca^2+^ influx through plasma membrane calcium channels(75)) are not affected by SLC29A3 (**Fig. 6E and Supp. videos**). Our data indicate that specifically TRPML1-dependent Ca^2+^ efflux from bacteria phagolysosomes, which accumulate nucleosides from bacteria degradation(76), is impaired in SLC29A3^-/-^ BMDCs.

**Figure 6.**
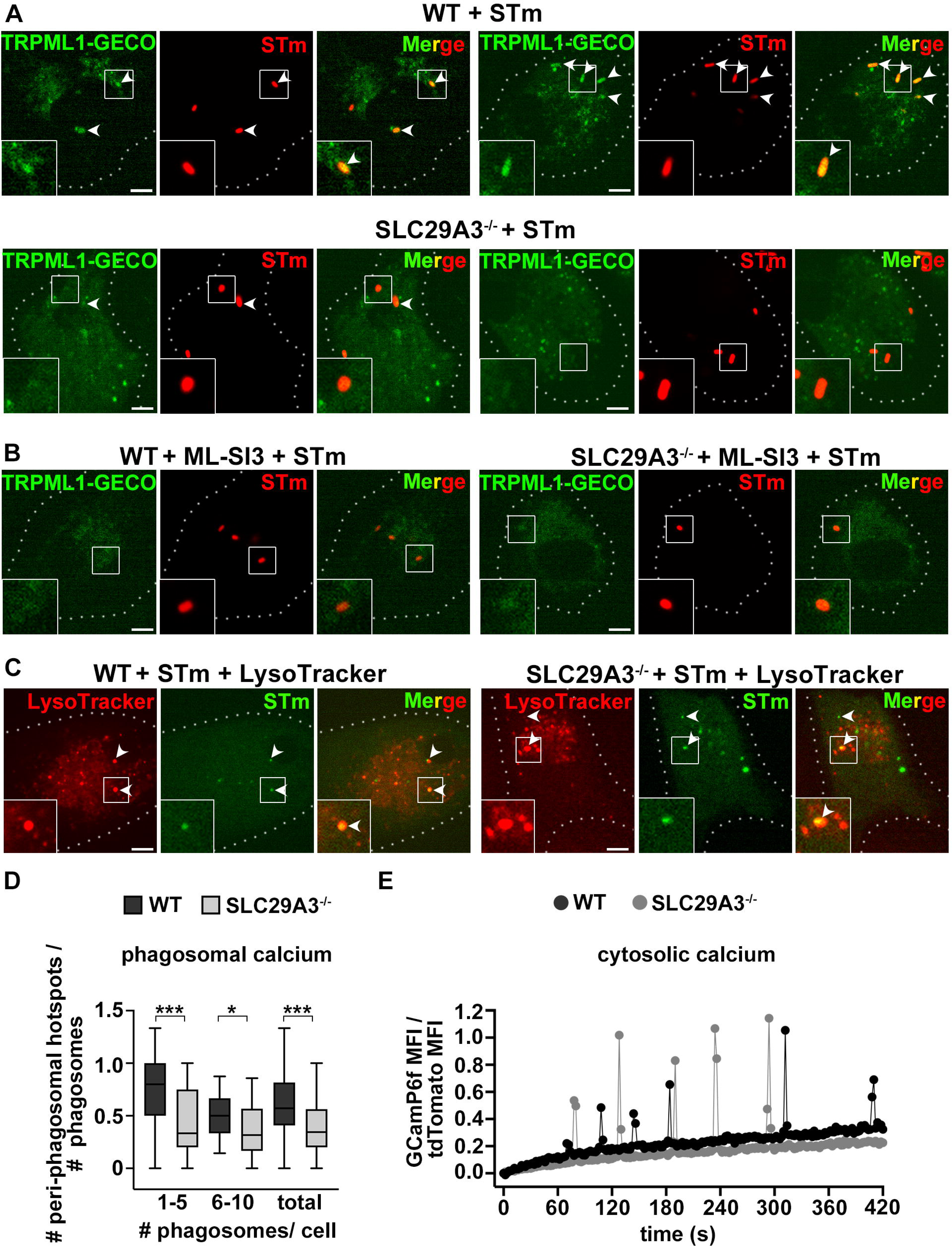
SLC29A3 promotes the activity of the lysosomal Ca^2+^ channel TRPML1. WT and SLC29A3^-/-^ BMDCs expressing TRPML1-GECO (A, B, D) or Salsa6f (E) or non-transduced (C) were stimulated with STm-mCherry at MOI=30 for 2 h (A), or pre-treated with ML-SI3 and stimulated with STm-mCherry at MOI=30 for 2 h (B), or pre-treated with LysoTracker red and stimulated with CFSE-labeled STm at MOI=30 (C), or stimulated with LPS-coated beads for 10 m (E), and visualized by live-cell imaging. **A.** Representative images showing TRPML1-GECO fluorescence (“hotspots”), indicative of calcium sensing. Arrows indicate peri-phagosomal “hotspots” on phagocytosed bacteria. **B.** Representative images showing absence of TRPML1-GECO activity (“hotspots”) in cells pre-treated with the TRPML1 inhibitor ML-SI3. **C.** Representative images showing CFSE-labeled STm colocalizing with or surrounded by lysosomes. **D.** Periphagosomal hotspots were quantified and compared to the total number of bacteria containing phagosomes in three independent experiments. **E.** The ratio of GCaMP6f and tdTomato MFIs was calculated with ImageJ overtime to assess cytosolic Ca^2+^ fluxes in response to stimulation with LPS-coated beads.

## Discussion

Dysfunctions in several cell types have been described as contributing to the various symptoms associated with H syndrome and SLC29A3 deficiency in mice(12-15). Particularly, histiocytosis – the accumulation of dysfunctional phagocytes – has been shown to contribute significantly to hyperinflammation in H syndrome and other types of histiocytic disorders(15, 77, 78). However, the mechanisms underlying both immunodeficiency and hyperinflammation in H syndrome are incompletely understood(3, 79). In addition, the contribution of DCs, key phagocytes in orchestrating both innate and adaptive immune responses and the main Ag presenting cells, has been previously unappreciated. Indeed, DCs participate in the initiation of anti-microbial pro-inflammatory responses and in the subsequent activation of T cell-mediated adaptive immune responses(17, 80). This dual behavior underscores DCs’ contribution to various inflammatory disorders(81).

Here we show that the lysosomal nucleoside transporter SLC29A3 is recruited to DC phagosomes after phagocytosis and supports their mild acidification to promote optimal phagosomal signaling and Ag MHC-II presentation to CD4+T cells (**Fig. 7**). We found that phagosomes in splenic DCs from SLC29A3^-/-^ mice bear significantly higher pH compared to WT, likely due to the accumulation of nucleosides and nucleobases in the absence of SLC29A3(13). This alkaline environment may impair phagosomal PRR signaling by limiting the availability of microbial degradation products or patterns, reducing the activation of PRRs in phagolysosomes(38, 82). In consequence, the nuclear translocation of the NF-κB subunit p65, as well as the subsequent production of IL-6, IL-12 and CCL-22, were reduced in either BM-derived or splenic DCs, in response to STm. Our data suggest that immunodeficiency in H syndrome may be partly due to impaired PRR-dependent phagolysosomal signaling to anti-microbial cytokines. The inability to clear the microbial insult will likely lead to the accumulation of defective DCs, which may contribute to sustain inflammation(83). Notably, the defective cytokine and chemokine secretion in SLC29A3^-/-^ BMDCs was rescued by the expression of wild-type human SLC29A3, but not the transport mutant SLC29A3^G437R^ (found in patients with H syndrome)(9), suggesting that transport activity is required for SLC29A3 role in anti-microbial signaling, at least partly by preventing the phagosomal accumulation of nucleobases. Interestingly, while IL-6 and IL-12 are pro-inflammatory cytokines, CCL22 coordinates contact between DCs and T-regs(84), suggesting that SLC29A3 deficiency might impair T-reg activation, further contributing to hyperinflammation. This possible scenario certainly warrants future investigations.

**Figure 7.**
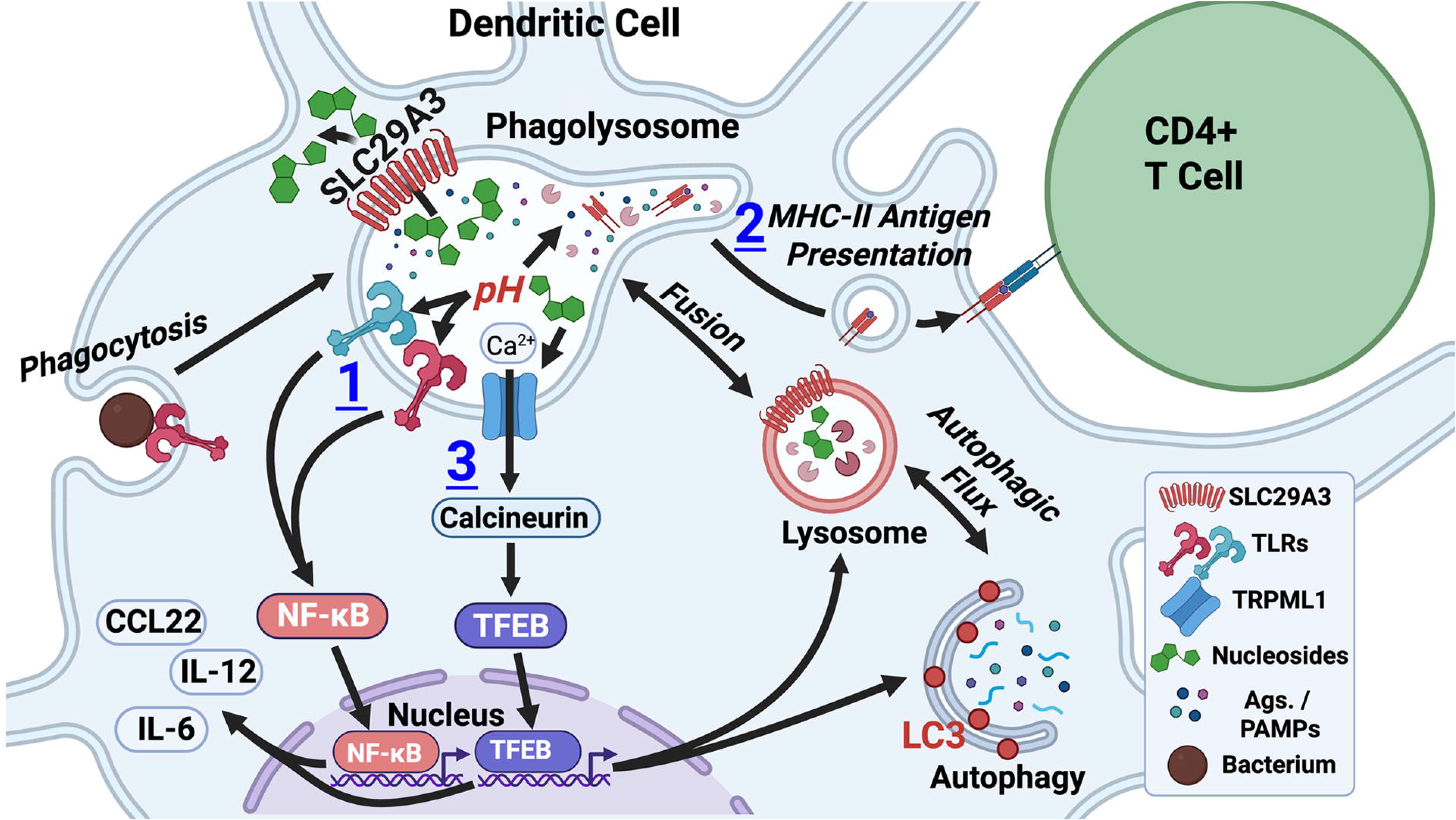
Schematics of the signaling pathways controlled by SLC29A3 in dendritic cells. After bacteria phagocytosis, lysosomal SLC29A3 is recruited to phagosomes during phagosome maturation. In turn, SLC29A3-mediated nucleoside export into the cytosol maintains intraphagosomal nucleoside and pH homeostasis. **(1)** SLC29A3 supports anti-bacterial phagosomal signaling to the production of cytokines and chemokines by sustaining phagosomal acidification and the generation of PAMPs for activation of phagosomal TLRs. **(2)** SLC29A3 promotes the formation of phagosomal tubules and MHC-II presentation of processed Ags. to CD4^+^ T cells to induce adaptive immune responses. **(3)** SLC29A3 promotes the activity of the phagolysosomal Ca^2+^ channel TRPML1, by preventing phagosomal alkalinization and adenosine accumulation. TRPML1 is triggered by phagocytosis and leads to the activation of calcineurin, which dephosphorylates the transcription factor TFEB, enabling TFEB nuclear shuttling. In the nucleus, TFEB induces the transcription of a gene network that promotes autophagy, and lysosome biogenesis and function, to control inflammation(45). TFEB also supports Ag. MHC-II presentation and anti-microbial gene programs (48, 66). *Figure created using Biorender.com through the TJU library portal*.

In addition to NF-κB-dependent pro-inflammatory signaling, TFEB has also been shown to promote the transcription of cytokines, chemokines and anti-microbial programs, besides autophagy(47, 48, 85). Indeed, TFEB is the master regulator of lysosomal function and autophagy and promotes lysosomal acidification and Ag MHC-II presentation(45, 66, 86, 87). In steady-state conditions, TFEB remains phosphorylated in the cytosol, which is mediated by mTORC1, a negative regulator of autophagy(46, 67-69). After phagocytosis, phagolysosomal Ca^2+^ efflux, mainly through the lysosomal channel TRPML1, promotes calcineurin-dependent TFEB dephosphorylation, driving TFEB to the nucleus and, therefore, promoting the activation of anti-microbial and autophagy programs(47). Notably, adenosine accumulation and alkalinization inhibit TRPML1(49, 54). Additionally, reduced cytosolic adenosine may inhibit AMP-activated protein kinase (AMPK), leading to mTORC1 activation(88, 89). Indeed, previous studies in hematopoietic stem cells have shown that adenosine accumulation in lysosomes due to SLC29A3 deficiency inhibits AMPK, activating mTORC1 and inhibiting autophagy(12). However, whether SLC29A3 may also promote autophagy by activating TRPML1 and TFEB after phagocytosis in DCs to boost anti-bacterial responses and control inflammation was previously unknown. We show that SLC29A3-deficiency in BMDCs causes decreased TFEB nuclear translocation and induction of autophagy after bacteria phagocytosis. As we and others have shown that autophagy is a negative regulator of inflammation(44, 90, 91), our findings suggest that defects in TFEB activation and autophagy may partly contribute to hyperinflammation in H syndrome and SLC29A3-deficiency in mice. Our data also show that SLC29A3-deficient BMDCs have reduced TRPML1 activation after bacteria phagocytosis, uncovering a previously unappreciated mechanism by which SLC29A3-deficiency may lead to hyperinflammation.

Both PRR and TFEB activation are known to promote Ag presentation on MHC-II molecules(21, 29, 58, 66). We show herein that while the expression of MHC-II molecules is similar between WT and SLC29A3^-/-^ splenic DCs, the presentation of the model Ag OVA, in soluble or particulate form, on MHC-II molecules to CD4+ T cells is impaired in SLC29A3^-/-^ splenic DCs. Given the striking defect in Ag MHC-II presentation, our data suggest that both the defective PRR stimulation and TFEB activation are contributing to this phenotype. In addition, our results show that the formation of phagosomal tubules, which we have found to promote MHC-II presentation by a TLR-dependent mechanism(58), is impaired in the absence of SLC29A3, suggesting that this phenotype could also contribute to the reduced Ag presentation. Solute carriers may also modulate membrane tension and dynamics(92). Indeed, SLC15A4 promotes lysosomal tubulation(37) and absence of SLC29A3 results in enlarged lysosomes(13). Further studies will be required to investigate the SLC29A3-dependent mechanisms driving phagolysosomal tubulation.

Activation of TLR7/8 by guanosine accumulation in SLC29A3 deficiency was shown to drive histiocytosis in humans and mice in the absence of microbial stimulation(15, 77). Our findings provide new insights in the context of bacteria phagocytosis and shed light on the previously unrecognized contribution of DCs to SLC29A3-deficiency. Our RNA seq analyses show differences in numerous transcriptional pathways in SLC29A3^-/-^ splenic DCs compared to WT in response to bacterial stimulation, highlighting that differences in phagosomal signaling appear to lead to large scale transcriptional changes. Overall, our findings support that SLC29A3 loss-of-function mutations may cause immunodeficiency and hyperinflammation by impairing both PRR signaling and TFEB activation upon microbial challenge. Defective but sustained cytokine and chemokine secretion by DCs (due to reduced autophagy and therefore impaired control of the inflammatory response), as well as defective T cell activation, likely prevents microbial clearance and contributes to the accumulation of immune cells. This scenario may sustain inflammation by resulting in an overall increase of released cytokines(83). As current strategies for the treatment of H syndrome include anti-inflammatory agents(77, 93), our findings suggest that modulation of autophagy or TFEB activity could also be considered to control inflammation in H syndrome, as proposed for other lysosomal storage disorders(94, 95).

## Materials and Methods

### Mice

C57BL/6 wild-type (WT) mice were originally purchased from The Jackson laboratories (Bar Harbor, ME). SLC29A3^-/-^ (C57BL/6J-*Slc29a3^em1(MPC)^*) mice were purchased from the Mutant Mouse Resource & Research Centers at The University of California, Davis. Sex- and age-matched mice between 8 and 12 weeks of age were used in all experiments.

### Ethics statement

Mice were bred under pathogen-free conditions at Thomas Jefferson University (TJU) and were euthanized by CO_2_ narcosis following the guidelines of the American Veterinary Medical Association Guidelines on Euthanasia. All performed animal studies were in compliance with regulations set by Public Health Service Policy on the Humane Care and Use of Laboratory Animals, the National Research Council’s Guide for the Care and Use of Laboratory Animals, the National Institutes of Health Office of Laboratory Animal Welfare, the American Veterinary Medical Association Guidelines on Euthanasia, and the guidelines of the Institutional Animal Care and Use Committee of TJU. All protocols used in this study were approved by the Institutional Animal Care and Use Committee at TJU (protocol #21-04-368).

### Reagents

LPS was purchased from InvivoGen (San Diego, CA), chlorophenol red-β-D-galactopyranoside (CPRG), bovine serum albumin (BSA) and ovalbumin (OVA) were from Sigma, OVA peptide (323-339) was purchased from InvivoGen, TxR-conjugated or Alexa Fluor 488 OVA, carboxyfluorescein succinimidyl ester (CFSE), BAPTA-AM and LysoTracker Red were from Invitrogen (Thermo Fisher Scientific) and ML-SI3 was from Selleck Chemicals, (Houston, TX). Rabbit anti-mouse p65 (D14E12), rabbit anti-mouse c-Jun (60A8), rabbit anti-mouse p38 (polyclonal), rabbit anti-mouse phospho-p38_T180/Y182_ (3D7), rabbit anti-mouse mTOR (7C10), rabbit anti-mouse phospho-mTOR_S2481_ (polyclonal), rabbit anti-mouse S6 (54D2), rabbit anti-mouse phospho-S6_S235/236_ (D57.2.2E), rabbit anti-mouse 4EBP1 (53H11), rabbit anti-mouse phosphor-4EBP1_T37/46_ (236B4), rabbit anti-mouse TLR7 (E4J3Z), rabbit anti-mouse GAPDH (14C10), rabbit anti-mouse histone H3 (D1H2), and rabbit anti-human TFEB (D207D) were purchased from Cell Signaling Technology (Danvers, MA). Mouse anti-β actin was from Sigma. Rabbit anti-mouse TFEB (polyclonal) was purchased from Bethyl Laboratories (Montgomery, TX). Rabbit anti-mouse LC3 (ab48394, polyclonal) and mouse anti-mouse p62 (ab56416) were purchased from Abcam Inc. (Waltham, MA). All secondary antibodies were from Jackson Immunoresearch (West Grove, PA) or Invitrogen. Allophycocyanin (APC)- and phycoerythrin (PE)-conjugated hamster anti-mouse CD11c (HL3), APC-conjugated rat anti-mouse CD11b (M1/70), FITC-conjugated mouse anti-mouse I-A^b^ (MHC-II; AF6-120.1), PE-conjugated mouse anti-mouse H2-kb (MHC-I; AF6-88.5), (PE-conjugated rat anti-mouse CD40 (3/23), FITC-conjugated rat anti-mouse CD86 (GL1), FITC-conjugated rat anti-mouse SIRPα (P84), FITC-conjugated rat anti-mouse CD4 (GK1.4), PE-conjugated rat anti-mouse CD8α (53-6.7), and FITC-conjugated rat anti-mouse B220 were purchased from BD Biosciences (San Diego, CA). Mouse APC-conjugated anti-mouse I-A^b^ (AF6-120.1) and rat PerCP-Cy5.5-conjugated anti-mouse F4/80 (BM8) were from Invitrogen.

### Cells and Cell Culture

Bone marrow cells were isolated from the femurs and tibiae of 8-16 week-old mice and cultured for 7-9 d in RPMI-1640 (Gibco, Thermo Fisher Scientific, Waltham, PA) media supplemented with 10% low-endotoxin FBS (HyClone, Cytiva, Logan, UT), 100 U/ml penicillin, 100 μg/ml streptomycin, 2 mM L-glutamine, 50 μM β-mercaptoethanol, and 30% granulocyte-macrophage colony stimulating factor (GM-CSF) containing media from J558L cells (a gift from Dr. R. Steinman’s former laboratory, Rockefeller University, NY) for differentiation to DCs as described(96).

For the isolation of splenocytes, spleens were digested with a solution of RPMI-1640 media containing 2 mg /mL collagenase D (Sigma-Aldrich, St. Louis, MO) and 10 μg/mL DNase I (Roche, Indianapolis, IN), at 37°C for 20 m(22). Single-cell suspensions were obtained by passing disaggregated tissue through 40 μm filters.

For the preparation of apoptotic splenocytes as a source of nucleosides, splenocytes were incubated at 47°C for 30 m and further incubated at 37°C for 4 h as described(15). Apoptosis was confirmed by annexin-V staining and flow cytometry.

Splenic DCs were isolated from single-cell suspensions of splenocytes by negative selection of B, T, and NK cells with a cocktail of biotin-conjugated antibodies (to CD90.2, CD45R, and CD49b) and anti-biotin microbeads (Miltenyi Biotec Inc., Auburn, CA), followed by positive selection with anti-CD11c (N418) microbeads (Miltenyi Biotec), and magnetic isolation on MACS columns (Miltenyi Biotec). Splenic DCs were phenotyped by flow cytometry and confirmed to be a mixed population of cDC1 (CD11c^+^ CD8^+^) and cDC2 (CD11c^+^ SIRPα^+^).

For experiments requiring large amounts of splenic DCs, 6-7 weeks old mice were injected subcutaneously in the abdomen with 5×10^6^ B16 cells producing Flt3-ligand (a gift from Dr. P. Roche, NIH), as recommended(97). Spleens were harvested around 15 days after injection, when the tumor mass was not bigger than 10 mm.

The B3Z and OT-IIZ hybridomas (which carry inducible NFAT-lacZ constructs)(65) (kindly provided by Dr. L. Eisenlohr, University of Pennsylvania, Philadelphia, PA), were cultured in RPM1 1640, 10% FBS, 100 U/ml penicillin, 100 μg/ml streptomycin, 2 mM L-glutamine, 50 μM β-mercaptoethanol, 1 mM sodium pyruvate, 1x MEM non-essential amino acids, 50 mg/ml geneticin or 50 mg/ml hygromycin.

### DNA retroviral constructs, retroviral production, and transduction of dendritic cells

Retroviral pMRX-IP TFEB-sfGFP was a gift from Eisuke Itakura (Addgene plasmid # 135402)^(98)^, TRPML1-G-GECO1.2-ERES was a gift from Antony Galione (Addgene plasmid # 207144)^(99)^, Salsa6f was a gift from Michael Cahalan (Addgene plasmid # 140188)^(75)^. TRPML1-G-GECO1.2-ERES and Salsa6f were subcloned into the retroviral backbone pMRX-IP at GenScript, (Piscataway, NJ). Human SLC29A3 was purchased from OriGene Technologies (Rockville, MD) and subcloned into the retroviral pMRX-IP-sfGFP downstream sfGFP at GenScript. SLC29A3 was mutated from G to A at position 1309 to generate the G437R mutation in the SLC29A3 protein (pMRX-IP-sfGFP-SLC29A3^G437R^) at GenScript.

Retrovirus was produced by transfection of the Platinum-E (Plat-E)(100) cell line using lipofectamine 2000 (Invitrogen). Supernatants from retrovirus-producing cells were harvested two days post-transfection. 3×10^6^ bone marrow cells were seeded per well on non-tissue culture treated 6-well plates. Two days after isolation, bone marrow cells were transduced by spinoculation with 3 ml of supernatant from transfected Plat-E cells with 8 μg/mL polybrene for 2 hours at 37°C. Following spinoculation, retrovirus-containing media were replaced with DC culture media. Puromycin (2 μg/ml) was added 3 d after infection, and cells were collected for experiments 3-4 d later.

### Bacterial culture

STm strains SL1344, SL1344 constitutively expressing mCherry and SL1344Δ*fljB*Δ*fliC* (STmΔfla) were kindly provided by Dr. Igor Brodsky (University of Pennsylvania, Philadelphia, PA). STmΔfla was grown overnight in 100 μg/ml streptomycin-containing LB medium at 37°C in an orbital shaker at 200 rpm. Bacteria concentration was determined by measuring optical density (OD) at 600 nm (1 OD_600_ equals 5×10^8^ bacteria/ ml). Bacteria were washed in PBS, then centrifuged at 13,500 xg and resuspended in DC media. For experiments with killed bacteria, STm was inoculated in LB medium in the morning and grown at 37°C in an orbital shaker at 200 rpm for 8 hours. Gentamicin (100 μg/ ml) was then added and the culture was incubated overnight to ensure bacterial death. The following day, bacteria concentration was measured as above, bacteria were washed in PBS, centrifuged at 13,500 xg and resuspended in DC media. Efficient bacterial killing was confirmed by plating bacteria on LB agar plates and incubating plates overnight.

### Bacterial stimulation

To measure cytokine secretion, 200,000 BMDCs or splenic DCs were seeded in DC culture media (without GM-CSF-containing supernatant) in triplicates in 96 well non-tissue culture-treated round bottom plates. DCs were pulsed with STmΔfla or gentamicin-killed STm at MOI=3 (for BMDCs) or MOI=10 (for splenic DCs) for 30 minutes. Gentamicin (100 μg/ml) was added to the cells treated with STmΔfla to kill extracellular bacteria after the pulse. Cells were chased for 3 hours at 37°C, centrifuged at 300xg for 5 minutes and supernatants were collected and tested for cytokine secretion using commercial ELISA kits (R&D Systems). Total RNA was isolated from cell pellets and subjected to RT-qPCR or RNA seq as described below.

For immunoblotting analyses, 2×10^6^ BMDCs were seeded on 35 mm tissue-cultured treated dishes. BMDCs were pulsed with STmΔfla at MOI=30 for 30 minutes. Gentamicin (100 μg/ml) was added after the pulse to kill extracellular bacteria. Cells were chased for 3 hours at 37°C, washed twice with RPMI-1640 and lysed in 200 μl Laemmli sample buffer.

### Real-time quantitative PCR (RT-qPCR)

Total RNA was isolated from BMDCs or splenic DCs using the RNeasy Mini kit (Qiagen, Germantown, MD). 1μg RNA was reversed transcribed to cDNA using TaqMan Reverse Transcription kit (Invitrogen). qPCR was performed in a StepOnePlus RT-PCR system using TaqMan master mix (Invitrogen) and TaqMan FAM probes (Applied Biosystems, Thermo Fisher Scientific) according to manufacturer’s instructions. TaqMan™ FAM probes specific for *il6*, *il12* and *ccl22* were Mm00446190_m1 (Il6), Mm01288989_m1 (Il12b) and Mm0436439_m1 (Il1b) and for the housekeeping genes β2-microglobulin (β2-m) and glyceraldehyde-3-phosphate-dehydrogenase (Gapdh) were Mm00437762_m1 (β2-m) and Mm99999915_g1 (Gapdh). PCR product formation was continuously monitored using the StepOnePlus™ Real-Time PCR System. Relative levels of mRNA were calculated with the ΔΔCt method(101), normalized to the average of housekeeping genes and represented as mRNA fold change. qPCR reactions were performed in triplicate. Experiments were independently performed three times.

### Live-cell imaging

BMDCs (7.5×10^5^) expressing retrovirally transduced GFP-SLC29A3 were seeded on poly-L-lysine coated glass-bottom 35 mm dishes (MatTek, Ashland, MA) on day 7-8 of culture. The following day, cells were pulsed with 3 μm Polybead microspheres (Warrington, PA) coated with TxR-conjugated OVA (1 mg/ml) and LPS (100 μg/ml) at MOI=10. Following the pulse, cells were washed with RPMI-1640 and the media was replaced with Leibovitz L-15 media (no phenol red; Gibco). Plates were imaged with a Nikon TiE dual channel spinning disk confocal microscope with a Hamamatsu Fusion sCMOS camera and equipped with an Okolab temperature and CO2-controlled chamber at the Sidney Kimmel Comprehensive Cancer Center (SKCCC) Bioimaging facility at TJU. Images were analyzed using ImageJ (NIH).

For the analyses of phagosomal tubulation, BMDCs were pre-loaded with apoptotic splenocytes (at a 20:1 ratio) for 2 days, seeded on poly-L-lysine coated glass-bottom 35 mm dishes and further pulsed with TxR-OVA/LPS beads. BMDCs were visualized 2 h after the pulse.

### Visualizing acidic organelles with LysoTracker Red

BMDCs (7.5×10^5^) were seeded on poly-L-lysine coated glass-bottom 35 mm dishes (MatTek), on day 7-8 of culture. The following day, 30-60 minutes before imaging, BMDC media was replaced with fresh media containing 50 nM LysoTracker Red (Invitrogen). Cells were incubated at 37°C for 30-60 minutes, media was replaced with fresh pre-warmed media, and cells were immediately imaged by live-cell imaging.

### Labeling of STm with CFSE

STm was collected at stationary phase and washed in PBS. 2×10^7^ bacteria were resuspended in 50 μL PBS pH 8.3 containing 100 μM CFSE and incubated at 37°C for 10 minutes as described(102). The reaction was stopped by the addition of 500 μL FCS. Bacteria were washed twice in PBS pH 7.2 containing 3% FBS, and once in PBS pH 7.2. Labeled bacteria were resuspended in 50 μL BMDC culture media before use.

### TRPML1-GECO imaging and calcium fluxes measurement

BMDCs (7.5×10^5^) expressing retrovirally transduced TRPML1-GECO or tdTomato-GCaMP6f (Salsa6f) were seeded on poly-L-lysine coated glass-bottom 35 mm dishes on day 7-8 of culture. The next day, for experiments with TRPML1-GECO, cells were pre-incubated with media containing 5 μM BAPTA-AM for 60 m before imaging to chelate cytosolic Ca^2+(73)^. Cells were pulsed with gentamicin-killed STm-mCherry at MOI=30 for 30 m. Following the pulse, cells were washed with pre-warmed Ca^2+^-free buffer (145 mM NaCl, 5 mM KCl, 3 mM MgCl2, 10 mM D-glucose, 1 mM EGTA, 20 mM HEPES) and media was replaced with Ca^2+^-free buffer containing 5 μM BAPTA-AM. For experiments to detect cytosolic calcium with Salsa6f, BMDCs were pre-incubated with apoptotic splenocytes 3 days before the experiment as described above. Cells were pulsed with STm at MOI=30 or LPS-coated polystyrene beads at MOI=10 with DC media without BAPTA-AM and visualized immediately after bacteria addition or 10 m after bead pulse. Cells were visualized by live cell imaging as described above. Frames were acquired every 2 seconds. Images were analyzed using ImageJ Time Series Analyzer plugin to calculate MFIs overtime.

### Immunostaining and flow cytometry

Cells were washed in flow buffer (PBS, 0.5% BSA, 2mM EDTA) and stained with fluorescently labeled antibodies in flow buffer on ice for 1 hour. Following immunostaining, cells were washed with flow buffer and analyzed by flow cytometry on a CytoFlex (Beckman-Coulter). Flow cytometry data were analyzed using FlowJo software (BD Biosciences).

### TFEB-GFP nuclear translocation by fluorescence microscopy

BMDCs (7.5×10^5^) expressing retrovirally transduced TFEB-GFP were seeded on poly-L-lysine coated glass-bottom 35 mm dishes on day 7-8 of culture. The following day, cells were pulsed with STmΔF at MOI=30 for 30 m. Gentamicin (100 μg/ ml) was added to kill extracellular bacteria. Following the pulse, cells were incubated at 37°C, chased for 180 m and imaged on a DMIL fluorescence microscope equipped with a K3C camera (Leica, Deerfield, IL). Images were analyzed using ImageJ (NIH).

### RNA sequencing (RNA seq) and analyses

Total RNA was isolated from BMDCs or splenic DCs using the RNeasy Micro kit (Qiagen, Germantown, MD). The quality and integrity of the RNA samples were assessed by the RNA integrity number (RIN) at the Genomics Core at TJU. Library preparation and RNA sequencing were performed at Novogene Corporation (Sacramento, CA) using Illumina NovaSeq 6000 PE150 sequencing platform and based on the mechanism of sequencing by synthesis. Original image data files were transformed to sequence reads (raw data) using Illumina Casava version 1.8. Raw reads were then filtered to remove adapter contamination, uncertain nucleotides and low quality nucleotides (base quality < 5) by Novogene. HISAT2 software was used for read alignment to the mouse reference genome. Further analyses were performed using the DEseq2 R package version 1.44.0(103). Principal component analysis (PCA) was performed on the 500 genes with the highest variance in the four data sets (WT untreated, WT treated, SLC29A3^-/-^ untreated, SLC29A3^-/-^ treated splenic DCs) after applying DESeq2 variance stabilizing transformation. Differential gene expression analysis was performed using the Wald test in DESeq2 and the p-values were adjusted using the false discovery rate correction method of Benjamini and Hochberg(104), as performed before(105). Significant genes were identified as having an adjusted p value < 0.01 and log2-fold change > 1. Enriched biological pathways were identified by performing gene ontology (GO) enrichment analysis on differentially expressed genes. Gene set enrichment analysis (GSEA) was performed against WT and SLC29A3^-/-^ groups using the fgsea R package(106). GO enrichment analysis was performed using the enrichGO function of the clusterProfiler R package(107). PCA and GO plots were visualized using the ggplot2 R package(108).

### Immunoblotting

Cells (200,000) were loaded per well in a 10 or 12-well 4-20% gradient polyacrylamide gel (Novex^TM^ Tris-Glycine, Invitrogen), transferred to PVDF membranes (Immobilon-P, Millipore, Burlington, MA) and analyzed using horseradish peroxidase-conjugated or fluorescently-labeled secondary antibodies (Jackson ImmunoResearch or Invitrogen), enhanced chemiluminescence (GE Healthcare, Pittsburgh, PA) and FluorChem R imaging system (ProteinSimple, Biotechne, San Jose, CA). Densitometric analyses of band intensities were performed using NIH Image J software, normalizing to control protein levels(109).

### Subcellular fractionation

Following stimulation with bacteria, BMDCs (5 x 10^6^) were incubated with 500 μL of cytosolic extraction buffer (1.5 mM MgCl_2_, 10 mM KCl, 10 mM HEPES, 1mM DTT, protease and phosphatase inhibitors cocktail) on ice for 10 m, detached from plates by scraping and transferred into microcentrifuge tubes. 0.05% (final concentration) NP-40 was then added, cells were incubated on ice for 3 m and centrifuged at 1,000 xg for 10 m. Supernatants were collected (cytosol-containing fraction). Pellets (nuclei-containing fractions) were resuspended in cytosolic extraction buffer with 0.1% NP-40, and incubated on ice for 3 m. Nuclei-containing fractions were centrifuged at 1,000 xg for 10 m, and resuspended in Laemmli sample buffer with 2-mercaptoethanol for immunoblotting.

### Phagosomal pH measurement

3 μm Polybead amino microspheres (Polysciences) were covalently coupled with FITC (Sigma; pH sensitive) and Alexa-Fluor (AF) 647 (Invitrogen; pH insensitive) for 4 hours at room temperature and washed as described(110). Briefly, BMDCs were pulsed with the FITC/AF647-coated beads at MOI=10 for 15 minutes, and extensively washed with PBS to remove extracellular beads. Cells were then incubated at 37°C for the indicated periods of time and immediately analyzed by flow cytometry (Cytoflex, Beckman Coulter) using a FITC/ AF647 gate selective for cells that had phagocytosed 1 bead. The ratio of the mean fluorescence intensity (MFI) emission between both dyes was determined. Values were compared with a standard curve obtained by resuspending the cells that had phagocytosed beads in CO_2_ independent medium (Invitrogen) at a fixed pH (ranging from pH 5 to pH 8) containing 0.1% Triton X-100.

### Proteomics on isolated phagosomes

WT BMDCs (40×10^6^) were incubated for 15 min with LPS-coated 3 μm magnetic beads (Dynabeads M-280 streptavidin, Invitrogen, NY) and then chased for 2 h. Cells were washed with PBS and seeded on a FBS cushion to separate non-internalized beads, as described(110) separated with disrupted with a syringe (22G needle) in 4 ml homogenization buffer (8% sucrose in PBS, 3 mM imidazole, 1 mM DTT and protease inhibitor cocktail), and phagosomes were isolated from the post-nuclear supernatant using a magnet, as we described before(63). Phagosomal fractions were characterized as shown in our previous publication(63), and subjected to mass spectrometry at the Children’s Hospital of Philadelphia Proteomics Core. Samples were run into an SDS PAGE gel, stained, excised and hydrolyzed with trypsin and LysC. The resulting peptides were de-salted, dried by vacuum centrifugation and reconstituted in 0.1% trifluoroacetic acid containing indexed retention time peptides (Biognosys Schlieren, Switzerland).

Peptides were analyzed on a Q-Exactive HF mass spectrometer (Thermo Fisher) coupled with an Ultimate 3000 nano UPLC system and an EasySpray source using data dependent acquisition (DDA), analyzed in Proteome Discoverer (Thermo Fisher) and Scaffold 5 software (Portland, OR). Peptide identifications were accepted if they could be established at greater than 0.0% probability by the Scaffold Local FDR algorithm. Protein identifications were accepted if they could be established at greater than 99.0% probability and contained at least 1 identified peptide. Protein probabilities were assigned by the Protein Prophet algorithm at the Proteomics Core(111).

### Antigen presentation

Splenic DCs (100.000) were pulsed with ovalbumin (OVA)/ bovine serum albumin (BSA)-coated beads, soluble OVA or OVA peptide ISQAVHAAHAEINEAGR (OVA_323–339_) at the indicated concentrations for 3 h. Cells were then co-cultured with 100,000 CD4+ OT-IIZ cells for 16 h at 37°C, as described before(112). Supernatants were collected for IL-2 detection by ELISA. Cell pellets were incubated with the β-galactosidase substrate CPRG in LacZ buffer (9 mM MgCl2, 0.125% NP40, 100 mM β-mercaptoethanol in PBS), for 4 h at 37°C. Optical density was measured at 590 nm with an Infinite F50 plate reader (Tecan US, Inc, Morrisville, NC).

### Statistical analyses

Statistical analyses were performed using Microsoft Excel and GraphPad Prism. Statistical significance was determined using the unpaired Student’s t-test after normality assessment. All experiments were performed independently a minimum of three times.

## Acknowledgments

We thank Stefan Feske (Ion Channels and Transporters in Immunity Research program, NYU) and Paul Roche (NIH, NCI) for critical reading of the manuscript, Paul Roche, Lawrence Eisenlohr, Igor Brodsky (University of Pennsylvania) and the former Ralph Steinman laboratory for the generous gifts of reagents, Michael S. Marks (University of Pennsylvania) for advice with the phagosomal proteomic analyses, Stefan Feske and Gaetan Barbet (Rutgers University) for advice with calcium flux measurements, Jason Hill and James Muller at the SKCCC BioImaging facility at TJU, Lynn Spruce at the Proteomics core at the Children’s Hospital of Philadelphia-Penn and Joan Kupper at the SKCCC Genomics Core at TJU for expert technical assistance, the SKCCC Flow Cytometry Core and the Office of Animal Resources at TJU.

This work was supported by NIH Grant R01 AI137173 (to C.L.-H. and A.R.M.), NIH-T32 AI134646 (to D.J.N.), and by the Intramural Research Program of the NIH, National Heart, Lung, and Blood Institute (NHLBI) ZIA HL006151 (to R.P. and J.A.M.). The BioImaging Shared Resource of the SKCCC is supported by NCI 5 P30 CA-56036. Fig. 7 was created using Biorender.com through the TJU library portal.

## Author Contributions

Daniel J. Netting designed and performed research, analyzed data, prepared figures and drafted the manuscript. Cynthia López-Haber designed and performed research and analyzed data. Zachary Hutchins analyzed RNA seq data and contributed to the preparation of the RNA seq figure. José A. Martina and Rosa Puertollano designed research and analyzed data. Adriana R. Mantegazza conceived, designed and supervised research, prepared figures and wrote the manuscript. All authors reviewed and edited the manuscript.

