## Supplementary figures and images for "The lysosomal carrier SLC29A3 supports anti-bacterial signaling and promotes autophagy by activating TRPML1 in mouse dendritic cells"

### Supplementary Figure 1

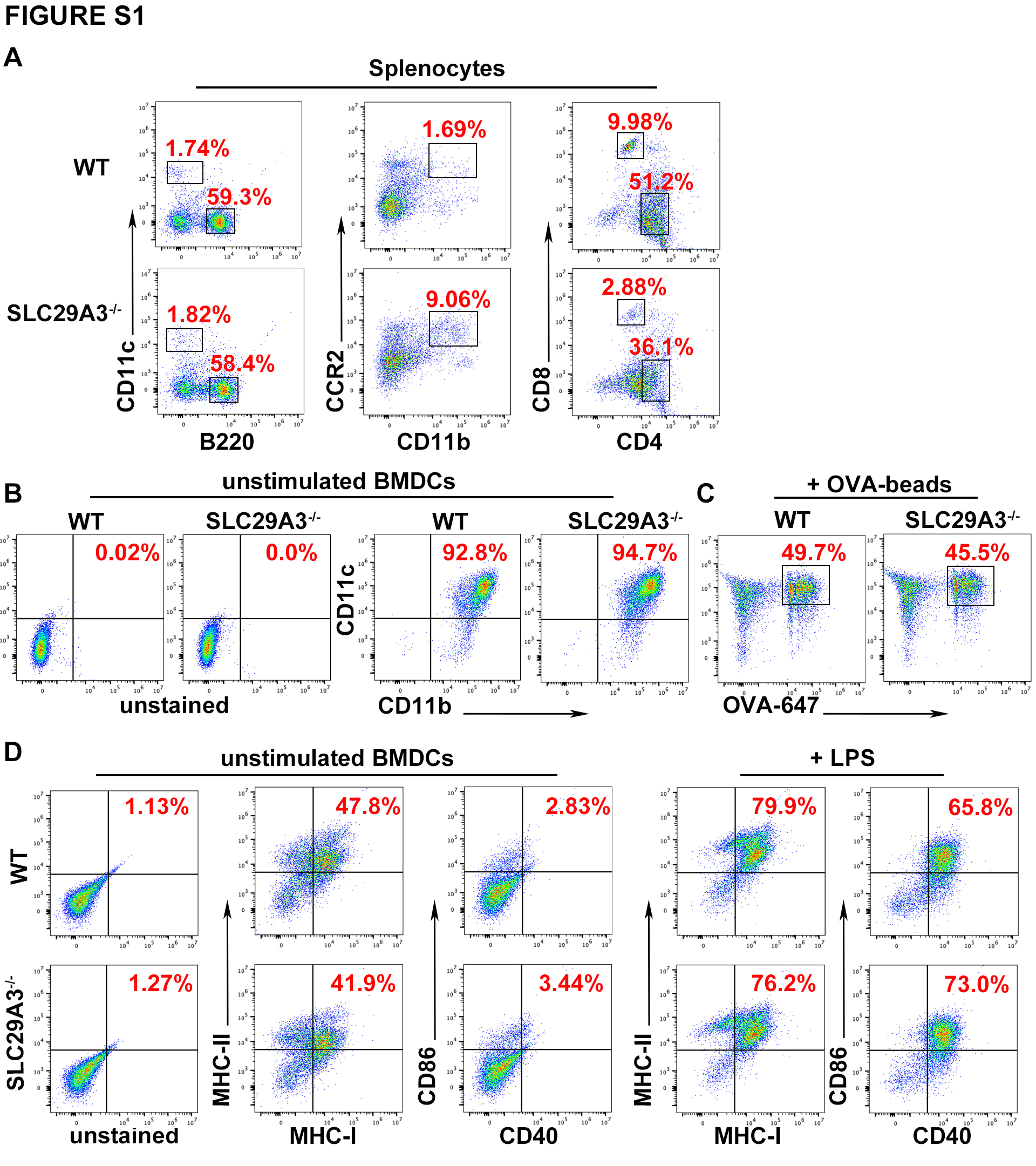

### Supplementary Figure 2

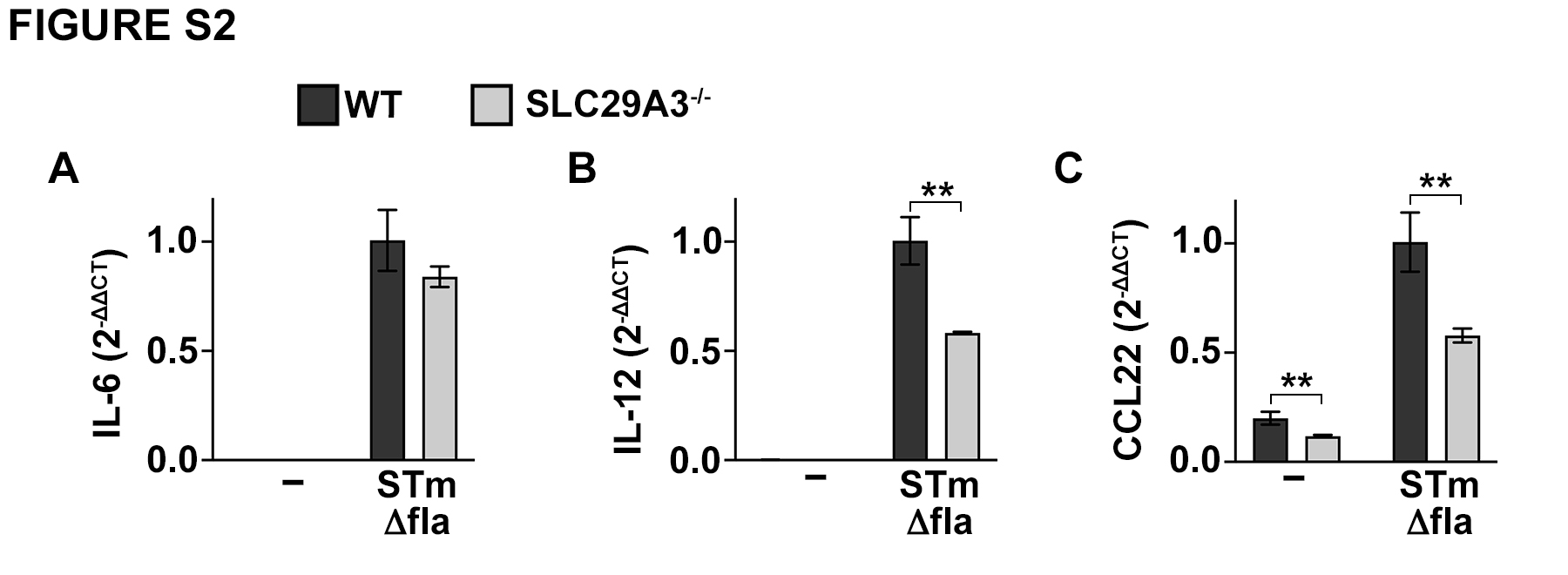

### Supplementary Figure 3

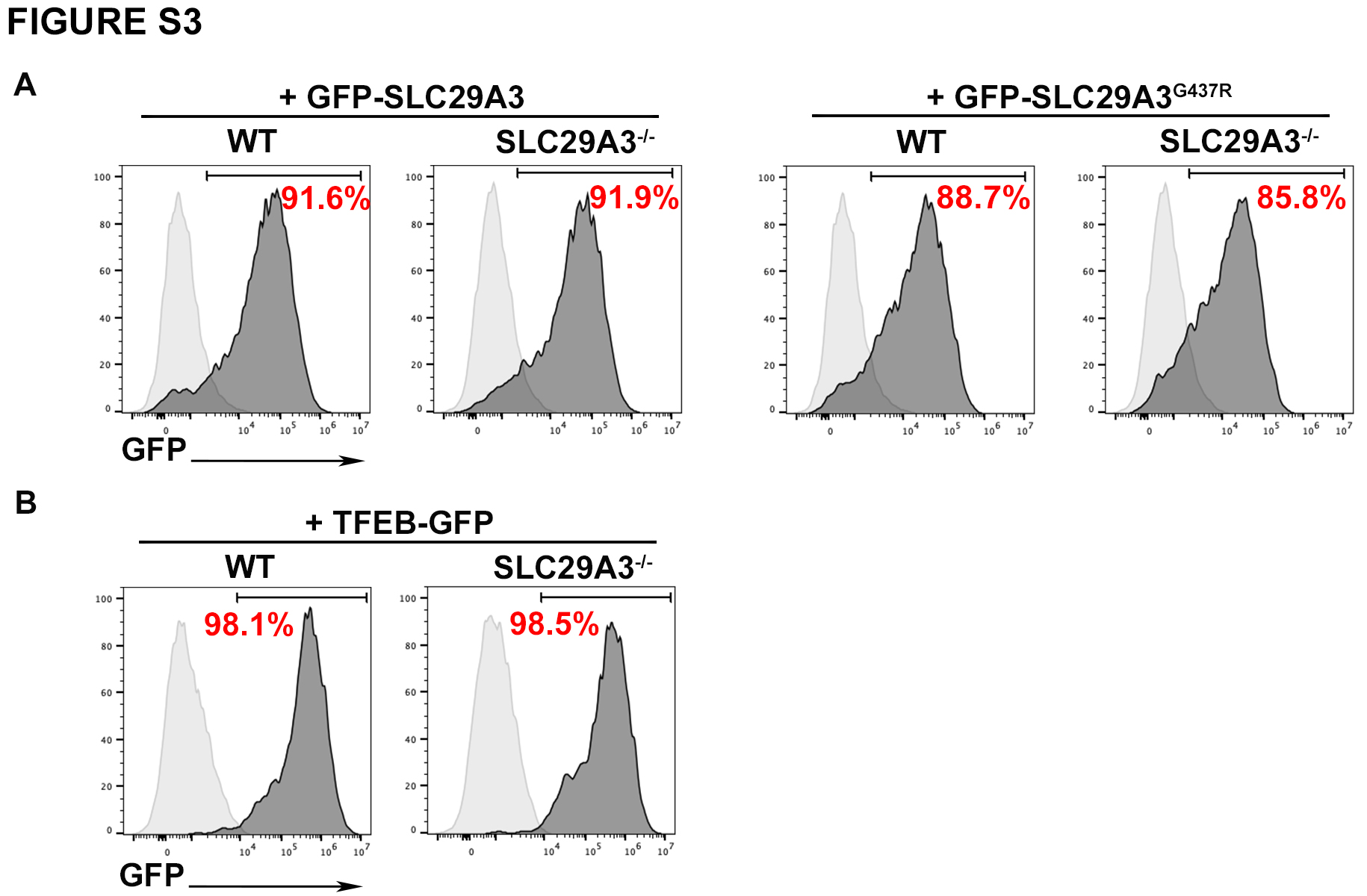

### Supplementary Figure 4

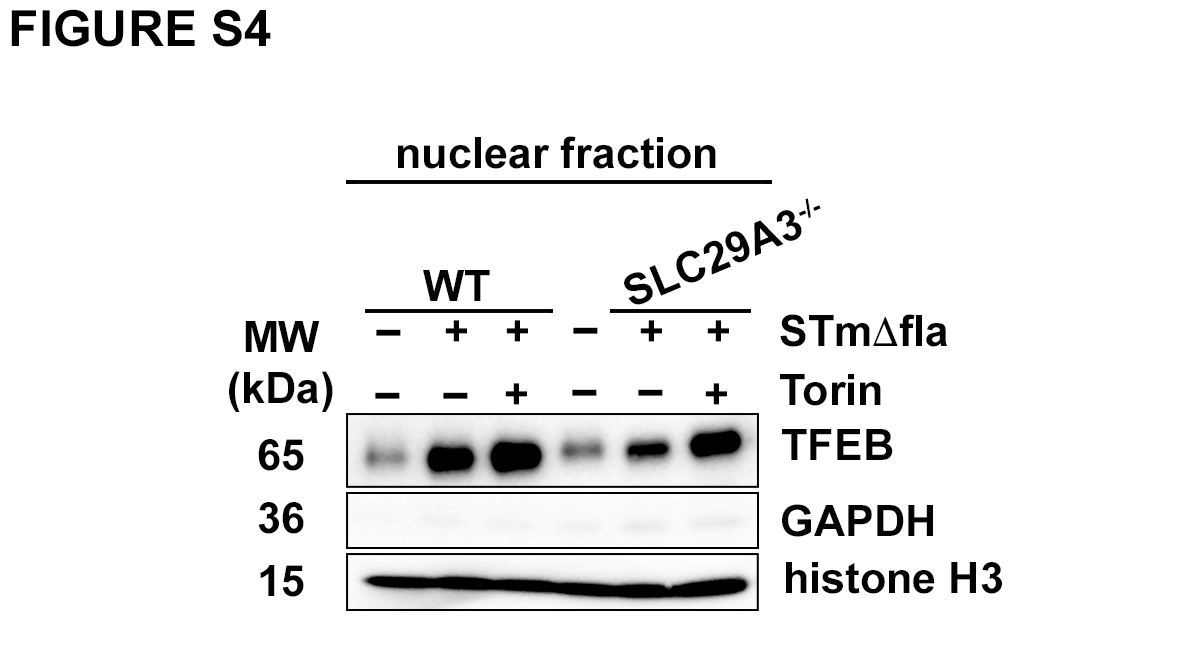

### Supplementary Figure 5

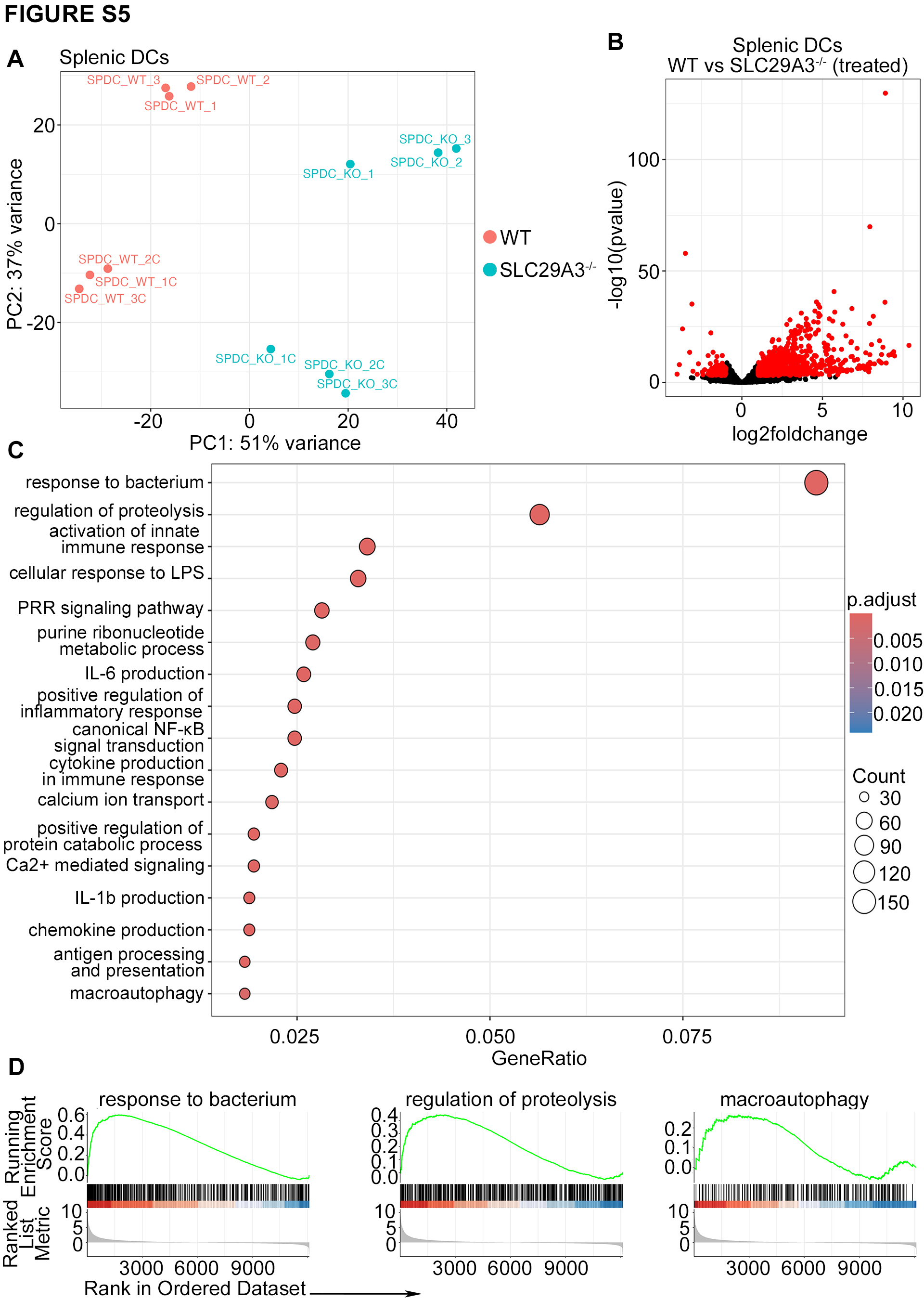
